# A discontinuity in motion perception during fixational drift

**DOI:** 10.1101/2025.06.02.657265

**Authors:** Josephine C. D’Angelo, Pavan Tiruveedhula, Raymond J. Weber, David W. Arathorn, Jorge Otero-Millan, Austin Roorda

## Abstract

The human visual system is tasked with perceiving stable and moving objects despite ever-present eye movements. Normally, our visual system performs this task exceptionally well; indeed, under conditions with frames of reference, our ability to detect relative motion exceeds the sampling limits of foveal cones. However, during fixational drift, if an image is programmed to move in a direction consistent with retinal slip, little to no motion is perceived, even if this motion is amplified. We asked: Would a stimulus moving in a direction consistent with retinal slip, but with a smaller magnitude across the retina, also appear relatively stable? We used an adaptive optics scanning light ophthalmoscope to deliver stimuli that moved contingent to retinal motion and measured subjects’ perceived motion, under conditions with world-fixed background content. We also tested under conditions with background content closer and farther from the stimuli. We found a sharp discontinuity in motion perception. Stimuli moving in a direction consistent with retinal slip, no matter how small, appear to have relatively little to no motion; while, stimuli moving in the same direction as eye motion appear to be moving. Displacing background content to greater than 4° from the stimuli diminishes the effects of this phenomenon.

## Introduction

Our eyes are never stable, even when fixating on a small target they make continuous miniature movements (Ratliff & Riggs, 1950). Despite this constant self-motion, the human visual system in general can properly perceive stable objects as stable and can detect moving objects with a level of precision that surpasses the sampling limits of the photoreceptors (Westheimer, 1975). To achieve this, the visual system requires world-fixed visual content which serves as a frame of reference (Legge & Campbell, 1981).

Yet, there exists a paradoxical exception: an image moving in a direction consistent with the direction of retinal slip appears relatively stable as long as the visual scene is filled with world-fixed retinal image background content (Arathorn, Stevenson, Yang, Tiruveedhula, & Roorda, 2013; D’Angelo, Tiruveedhula, Weber, Arathorn, & Roorda, 2024; Riggs, Ratliff, Cornsweet, & Cornsweet, 1953); and when all visual content is removed, this same image appears to be moving with a significantly higher magnitude of motion (D’Angelo et al., 2024). This phenomenon suggests that information from the retinal image informs the visual system about its direction of eye motion, and renders everything in the perceived image that is moving in a direction consistent with retinal slip as being relatively stable.

This “illusion of relative stability” suggests that there is a discontinuity in motion perception during drift, where direction is the critical parameter that governs how motion is perceived. Specifically, for images moving in a direction consistent with retinal slip, little to no motion is perceived, and for images moving in any other direction, the motion is readily detectable. In this present study, we assessed the sharpness of this discontinuity by measuring the perceived motion of stimuli that moved in a direction consistent with retinal slip with decreasing magnitudes, and of stimuli that moved in the same direction as eye motion. Next, we determined the extent to which the world-fixed retinal image background content drives this phenomenon by performing a second experiment in which we varied the proximity of the background content to the stimuli.

Our findings suggest a sharp discontinuity in motion perception during fixational drift. Even a stimulus moving in the same direction as eye motion but with a slower speed, and therefore slipping opposite to eye motion with 10% of the retinal motion of a world-fixed stimulus, appears relatively stable. We show that, even if a stimulus is moving on top of or very close to background edges, as long as it moves opposite to eye motion, it appears relatively stable. Thus, this phenomenon depends on the presence of retinal image background content within several degrees of the stimuli, and moving background content beyond ~4° diminishes the effect. This work builds upon our understanding of this paradoxical phenomenon that disrupts our hyperacute sensitivity to detect relative motion.

## Methods

### Subjects

Five subjects provided informed consent and participated in the experiments [subject number, age, sex]: [10003L, 58, M], [20237R, 27, F], [20255L, 24, M], [20256R, 23, F], and [20258R, 30, F]. One subject is a co-author on this paper and the other four subjects were experienced with psychophysical tasks. All procedures were approved by the Institutional Review Board at the University of California, Berkeley. Prior to the experiment, subjects applied one drop of 1% tropicamide and one drop of 2.5% phenylephrine hydrochloride to dilate and cycloplege.

### Adaptive Optics Scanning Light Ophthalmoscopy

Retinal imaging and stimulus delivery were performed using a multiwavelength AOSLO (Mozaffari, LaRocca, Jaedicke, Tiruveedhula, & Roorda, 2020; Roorda et al., 2002). Light from a supercontinuum laser (SuperK Extreme; NKT Photonics, Birkerod, Denmark) was split into 3 independent channels. The 940-nm wavelength was used to continuously measure the optical imperfections of the eye, which were corrected with a deformable mirror (DM97-08; ALPAO, Montbonnot-Saint-Martin, France), enabling near diffraction-limited resolution. The 840-nm wavelength imaged a 1.71° field of the retina. The 680-nm wavelength was modulated by an acousto-optic modulator (AOM) to deliver three independent stimuli (Fig. 1A). This was possible by updating the FPGA board to allow up to three retina-contingent stimuli, rather than one. This new feature was an upgrade from previous work (D’Angelo et al., 2024). The system had a vertical frame rate of 60 Hz and a fast horizontal scan rate of ~16 kHz, rendering 512 by 256 pixel video recordings. Each pixel subtended 0.2 by 0.4 arcmin of visual angle. The average powers of the 940-nm, 840-nm, and 680-nm lasers were 50.3 *µ*W, 110.1 *µ*W, and 3.53 *µ*W, which correspond to equivalent luminance values of 0.0031 cd/m^2^, 0.83 cd/m^2^, and 1,200 cd/m^2^ using methods described in Domdei et al. (2018).

**Figure 1:**
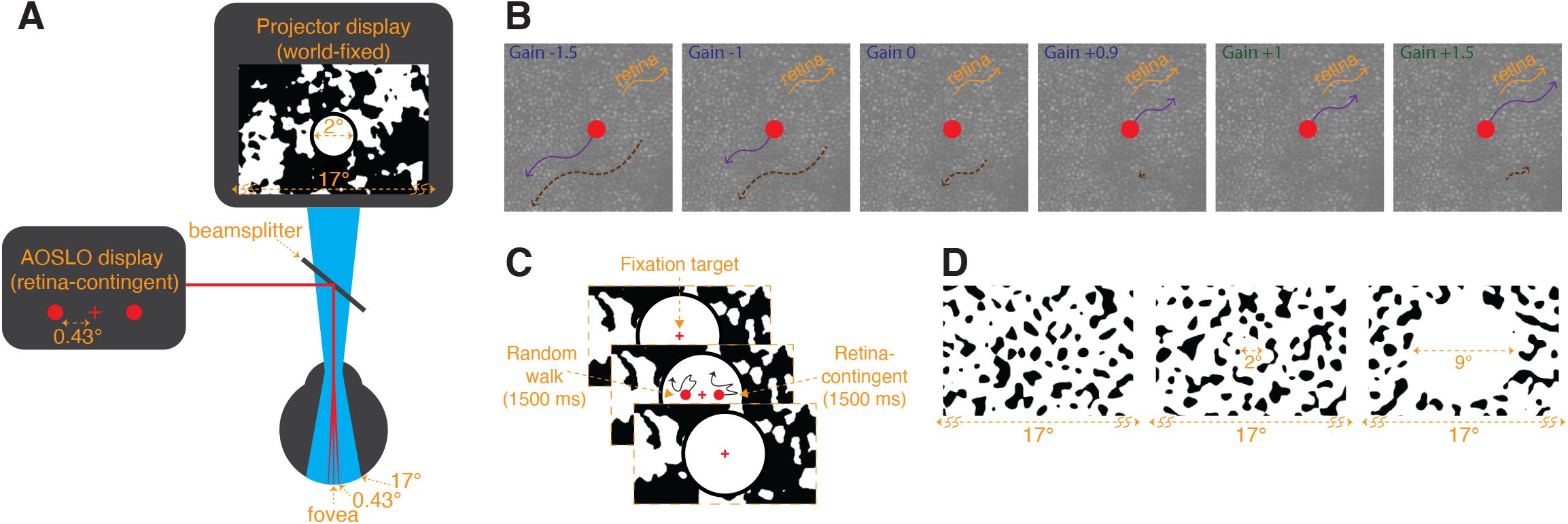
Display configuration. (A) The AOSLO drew three 680-nm stimuli: a central fixation cross, and two horizontally offset circular stimuli positioned 0.43° nasal and 0.43° temporal from the cross. Concurrently, over its 17° field of view, the projector displayed world-fixed patterns, dependent on the experiment. In Experiment 1, the background was filled with a blurred and binarized 1/f noise pattern with a 2°-diameter central white circle with a surrounding black ring that overlayed the 680-nm stimuli, shown in (A). In Experiment 2, the background was filled with a noise pattern that was generated by filtering white noise with a Gaussian bandpass filter at 0.5 cycles per degree (cpd) (0.5 cpd standard deviation), and binarizing the pattern, shown in (D). (B) Rules for the retina-contingent stimuli. Each panel shows examples of different retina-contingent stimuli undergoing identical retinal trajectories, indicated by the orange arrows. The purple arrows indicate the world trajectories of the retina-contingent stimuli. The dashed-brown arrows indicate the stimuli’s image motion across the retina. Gains *>* 1 move in the same direction as eye motion (orange and dashed-brown arrows point in the same direction) while Gains *<* 1 move opposite to eye motion (orange and dashed-brown arrows point in opposite directions). (C) Experimental sequence. Subjects fixated on a world-stationary cross and initiated a 1500-ms presentation of two stimuli that moved with different rules. The left stimulus moved in a random walk, independent of eye motion, and the right stimulus moved contingent to eye motion with a Gain that ranged from −1.5 to +1.5. (D) Experiment 2 background conditions. The patterns extended over the full 17° field of view (Left), or surrounded a 2°-diameter white circular opening (Middle), or surrounded a 9°-diameter white circular opening (Right). The AOSLO stimuli were presented at the center of the displays and the subject performed the same experimental protocol shown in (C). For the no-white-circle and 2°-diameter white circle conditions, the fixation cross remained on for the entire duration. For the 9°-diameter white circle condition, the fixation cross turned off during the 1500-ms presentation of the stimuli.

#### Real-time Eye Tracking

When imaging with an AOSLO, eye movements expand, compress, or distort the retinal images. To mitigate these effects, the system recovers and then corrects for the eye movements in real-time. This is achieved through the following: At the beginning of the experiment, the experimenter selects a reference frame. During the experiment, as the 512 by 16 pixel strips are acquired by the fast horizontal scanner, they are registered to the reference frame. The x and y offsets necessary for best registration are measures of the absolute eye movements during the recording.

#### Targeted Stimulus Delivery

By recording the eye motion in real-time, the AOSLO can perform targeted stimulus delivery. As the 840-nm laser images the retina and the system recovers the eye motion in real-time, an AOM arms the 680-nm laser to deliver stimuli that move contingent to eye motion (Arathorn et al., 2007). These stimuli are called “retina-contingent stimuli”. The use of narrow strips for registration combined with fast processing, enables a 2-ms lag between eye motion prediction and stimulus delivery (Stevenson & Roorda, 2005; Stevenson, Roorda, & Kumar, 2010; Yang, Arathorn, Tiruveedhula, Vogel, & Roorda, 2010).

A retina-contingent stimulus has a “Gain” which describes the stimulus’ world motion, that is contingent on the eye motion. The movements of the retina-contingent stimulus are equal to the eye movements multiplied by the Gain. The sign of the Gain indicates the direction the stimulus moves with respect to the eye movement. Gains *<* 0 move directly opposite to eye motion with amplified retinal slip. For example, the world motion of a Gain −1 stimulus is equal to −1 times the eye motion (Fig. 1B, panel 2, purple arrow) and its motion across the retina is twice that of a world-fixed stimulus (Fig. 1B, panel 2, dashed-brown arrow). Gain 0 describes a world-fixed stimulus, its world motion is equal to 0 times the eye motion (Fig. 1B, panel 3, no purple arrow) and the stimulus naturally slips consistent to drift motion (Fig. 1B, panel 3, dashed-brown arrow). Gain +1 describes a stabilized stimulus, its world motion is equal to 1 times the eye motion (Fig. 1B, panel 5, purple arrow) and therefore is locked in place on a region of the retina (Fig. 1B, panel 5, no dashed-brown arrow). Gains *>* 0 move faster than the eye motion and in the same direction. For example, the world motion of a Gain +1.5 is equal to +1.5 times the eye motion (Fig. 1B, panel 6, purple arrow) and its motion across the retina is half that of a world-fixed stimulus (Fig. 1B, panel 6, dashed-brown arrow).

Note that Gains greater than 0 but less than +1 move in a direction consistent with the direction of retinal slip, but with reduced retinal slip compared to a world-fixed stimulus. For example, if the eye moves to the right by 10 arcmin, a Gain +0.9 stimulus would move right by 9 arcmin with retinal slip equal to 1 arcmin. Although both move in a direction consistent with retinal slip, Gain +0.9 has only 10% the retinal slip of a world-fixed stimulus. The retinal slip of the Gains 0 and +0.9 are illustrated by dashed-brown arrows in Fig. 1B in Panels 3 and 4, respectively. Gain +1 has zero retinal slip, and is therefore the transition point: For all Gains *<* +1 the stimulus’ motion across the retina is in a direction directly opposite to eye motion, in a direction consistent with retinal slip; and for all Gains *>* +1, the stimulus’ motion across the retina is in the same direction as eye motion.

### Projector Display

A digital light projector (Lightcrafter DLP4500; Texas Instruments, Dallas, TX, USA) was used to display a 17° background that was coaligned with the AOSLO beams (Fig. 1A). The average luminance was approximately 540 cd/m^2^. Background patterns were programmed with MATLAB (MathWorks, Natick, MA, USA) using the Psychophysics Toolbox (Brainard, 1997; Kleiner et al., 2007; Pelli, 1997).

### Random Walk Analysis

Fixational drift is similar to a random walk and the magnitude of drift can be quantified by computing a diffusion constant (Clark, Intoy, Rucci, & Poletti, 2022; D’Angelo et al., 2024). The diffusion constant represents the amount the eye moves from its starting position as a function of time. Recent studies have found that drift motion is not truly random, instead having correlated or anti-correlated properties depending on the timescale (Engbert & Kliegl, 2004; Engbert, Mergenthaler, Sinn, & Pikovsky, 2011), which can be quantified by computing the parameter *α* (D’Angelo et al., 2024; Engbert & Kliegl, 2004; Engbert et al., 2011; Roberts, Wallis, & Breakspear, 2013; Mergenthaler & Engbert, 2007). We evaluated the mean square displacement (MSD) as a function of multiple time intervals (△T), to determine the magnitude of drift (D) and the extent to which the drift statistics deviate from Brownian motion (*α*), by fitting the model shown in Equation 1 to the empirical. The dimension (d) is equal to two because we recorded the x and y positions of the eye. As shown in Equation 2, the empirical MSD was computed over non-overlapping pairs, where N is equal to the total number of time steps (total number of frames) and ⌊ ⌋ represents the greatest integer function. We plotted the *log*_10_(MSD) as a function of the *log*_10_(△T). The slope of the line across these data points is *α* and the y-intercept is *log*_10_(diffusion constant). The MATLAB code and full description of these parameters and methods are reported in D’Angelo et al. (2024).

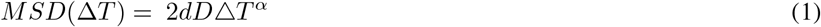

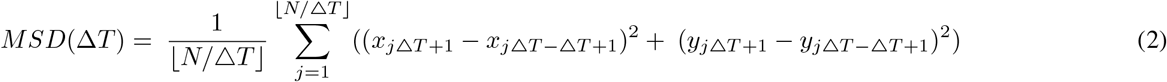

In this study, we use the terms “Brownian motion” to describe uncorrelated motion (*α* = 1), “persistence” to describe positively correlated motion (*α >* 1), and “antipersistence” to describe negatively correlated motion (*α <* 1). We performed this random walk analysis on the random walk stimulus’ motion (*D*_*PM*_ and *α*_*PM*_, where PM denotes perceived motion), the retina-contingent stimulus’ motion (*D*_*WM*_ and *α*_*WM*_, where WM denotes world motion), and the eye’s motion (*D*_*EM*_ and *α*_*EM*_, where EM denotes eye motion).

### Generating Random Walks and Random Jitter

Before the experiments, we generated 100s of 1500-ms duration random steps that were either cumulatively summed to produce random walks or concatenated to produce random jitter. The method to generate random walks is described more fully in D’Angelo et al. (2024). Briefly, step lengths in x and y were drawn from a normal distribution with standard deviations ranging from 0.05 to 1.6 arcmin per step in increments of 0.05 arcmin. We computed the average diffusion constant from the trajectories generated for each step length. This was achieved by first solving for the mean square displacement in Equation 2, where △X and △Y are equal to the step length, and then using this result in Equation 1 to solve for the diffusion constant. For each step length, we generated 100 random walks and then selected 10 paths of the 100 which had diffusion constants closest to the respective average diffusion constant. The *α* of the random walks was approximately equal to one. To program random jitter, step lengths in x and y were drawn from a normal distribution with the following standard deviations: 0.1, 0.2, 0.3, and 0.4 arcmin per step. These steps were non-cumulative, meaning that each step began from the same starting position. Therefore, the *α* of the random jitter was approximately equal to zero.

### Experimental Design

#### Set Up

Subjects’ aligned the pupil of one eye with the AOSLO beam and covered their fellow eye with an eye patch. Dental bite bars were used to immobilize the head. The bite bar apparatus was adjusted in the X, Y, and Z dimensions to position the eye’s entrance pupil at the exit pupil of the AOSLO and the projector to optimize wavefront correction performance and to optimize the image of the retina.

#### Experimental Procedure

The subject was presented three 680-nm stimuli through the AOSLO: a central fixation cross, and two horizontally offset circular stimuli positioned 0.43° nasal and 0.43° temporal from the cross. The fixation cross subtended 5 arcmin and the circular stimuli subtended 6 arcmin. Concurrently, the projector displayed a 17° background that was superimposed with the AOSLO rasters (Fig. 1A). In the first experiment, the background displayed a blurred and binarized 1/f noise pattern that changed after each presentation. A 2°-diameter white circle, with a 0.33°-wide black ring around it, was overlayed onto the AOSLO rasters, therefore canceling perception of the 840- and 940-nm rasters (Fig. 1A). The fixation cross remained on for the entire experiment.

In the second experiment, the background displayed a noise pattern that was generated by filtering white noise with a Gaussian bandpass filter at 0.5 cycles per degree (cpd) (0.5 cpd standard deviation), and then binarizing the pattern. We tested three background conditions: extending over the full 17° field of view (Fig.1D, Left), or with a 2°-diameter white circular opening (Fig.1D, Middle), or with a 9°-diameter white circular opening (Fig.1D, Right). Similar to Experiment 1, the patterns changed after each presentation. Under the no-white-circle condition, the pattern extended over the AOSLO rasters, and therefore, the 840- and 940-nm rasters were dimly visible in the regions where the pattern was black (Fig.1D, Left). For the no-white-circle and 2°-diameter white circle conditions, the fixation cross remained on for the entire experiment while for the 9°-diameter white circle condition, the fixation cross turned off during the 1500-ms presentation of the stimuli. For the 2°- and 9°-diameter white circle conditions, the subjects’ field of view was limited to only 17°. Using methods described in D’Angelo et al. (2024), we set up a white paper with an aperture in front of the display permitting only the AOSLO and projector beams to enter the eye. Notably, the aperture in our experiment was wider than the one used in D’Angelo et al. (2024) yielding a full 17° view of the projector and AOSLO displays. LEDs were positioned between the subject and the paper to illuminate the paper. Due to its close proximity to the eye, the natural blur of the aperture in the paper rendered the transition between the display and luminance-matched paper invisible. The different choices for the binarized patterns from Experiment 1 and 2 yielded no difference in the results.

#### Retina-Contingent Conditions

In the first experiment, we measured the perceived motion of multiple retina-contingent stimuli. The Gains of the stimuli ranged from −1.5 to +1.5 in increments of 0.25. We additionally tested Gains +0.85, +0.9 and +1.1 to measure closer to Gain +1, the transition point between stimuli that move opposite to eye motion and stimuli that move in the same direction as eye motion. A total of sixteen Gains were tested and the subjects performed at least three trials per Gain. Subjects performed at least six trials of Gain +1.25 and +0.75. Subjects performed nine trials of Gain +1, to mitigate the effects of uncertainty due to fading (except for 10003L, who only performed six trials of Gain +1). In the second experiment, we tested only Gains −1.5, 0, and +1.5 and subjects performed at least four trials per Gain (except for 20237R, who only performed three trials for Gain −1.5 of the 9°-diameter white circle condition).

#### Task

One Gain was tested per trial. In a single trial, the subject began by fixating on the 680-nm cross. Using a gamepad, the subject initiated a 1500-ms presentation of two horizontally offset circular 680-nm stimuli. The three 680-nm stimuli were programmed with different rules: the fixation cross remained stationary, the left stimulus moved in a pre-programmed random walk that was independent of eye motion, and the right stimulus moved contingent to the retina with a specific Gain (Fig. 1C). After attending to the circular stimuli, the subject adjusted the magnitude of motion of the left stimulus (random walk) until it matched that of the right stimulus (retina-contingent). The subject could initiate as many presentations as they wished while making their adjustments. When they reached a match, they were instructed to initiate at least three more presentations before submitting their response. The subjects were not given any specific criteria to make the match and in general expressed satisfaction in their ability to achieve one. After submitting the trial, the subject proceeded to the next trial which tested a new retina-contingent stimulus with a specific Gain. The Gains were presented in a pseudorandom order across multiple sessions. This experimental protocol applied to both Experiments 1 and 2.

### Eye and Stimulus Motion Traces

From each presentation of the experimental sequence, we recorded a two-second duration video of the retina. Additionally, the system drew three digital markers that indicated the exact positions of the three 680-nm stimuli on each frame of the video. After the experiment, we extracted absolute eye motion traces from the videos through an offline software called ReVAS, which uses a similar cross-correlation method described in Section **Real-time Eye Tracking** (Agaoglu, Sit, Wan, & Chung, 2018). Next, we used an offline algorithm to locate and record the positions of the digital markers in each frame. The eye motion and stimulus motion traces were used for the following:

#### Quality Control

First, we identified and removed videos where tracking failed. Tracking failure results in frames with no image captured or with poor image quality and ReVAS either fails to extract eye motion traces or produces fragmented traces. Tracking can fail for a variety of reasons: for example, if the subject blinks, saccades, or looks away from the fixation cross during the presentation. Dry tear film or a fallen eyelash can also disrupt the wavefront sensor and lead to tracking failures. These were usually recognizable to the subjects, who were instructed prior to the experiment to ignore presentations with poor tracking.

Second, we identified and removed videos where the tracking did not fail, but a microsaccade occurred during frames in which the stimuli were delivered. We kept a record of the number of videos per trial where a microsaccade occurred. Our goal was to measure the perceived motion during periods of fixational drift, and therefore including presentations with microsaccades would have confounded the analyses.

Third, we determined the number of videos where the tracking was poor but might not have been recognizable to the subject. This was achieved by comparing the retina-contingent stimulus motion trajectory to the eye motion trace. For every video, we computed the ideal retina-contingent stimulus motion trajectory computed from the eye motion trace and the specific Gain. For each frame, we computed the distances in x and y between the ideal retina-contingent position and the measured retina-contingent position. This distance represented the stimulus misdelivery, where greater distances indicate greater error in stimulus delivery. For each dimension, we took the standard deviation of the distances across all frames. If the standard deviation of the stimulus misdelivery in either the x or y dimension was greater than 0.9 arcmin, we removed the video from further analysis. We tallied the number of poor videos – videos that had standard deviations greater than 0.9 arcmin or that contained microsaccades − and if this number was greater than half the total number of videos in the trial, we dropped the trial and the subject was invited back to repeat the trial.

#### Quantifying the Stimulus Delivery Error

With the remaining traces, we quantified the magnitude of stimulus delivery error for each trial. Using the same method described in Section **Quality Control**, we computed the distances in x and y between the ideal retina-contingent position and the measured retina-contingent position. In a single trial, we computed the standard deviation of all distances in x and y across all frames from all videos. An example from one subject is shown in Fig. 2A, the red points indicate the standard deviation of error from a single trial and the black points indicate the average standard deviation of error across all trials. Note that this subject, 10003L, had the highest error out of the five subjects tested, with a maximum average standard deviation of error equal to ~0.48 arcmin, which is about the diameter of a foveal cone photoreceptor (Y. Wang et al., 2019). The results of all subjects can be found in Fig. A1.1 in Appendix 1.

**Figure 2:**
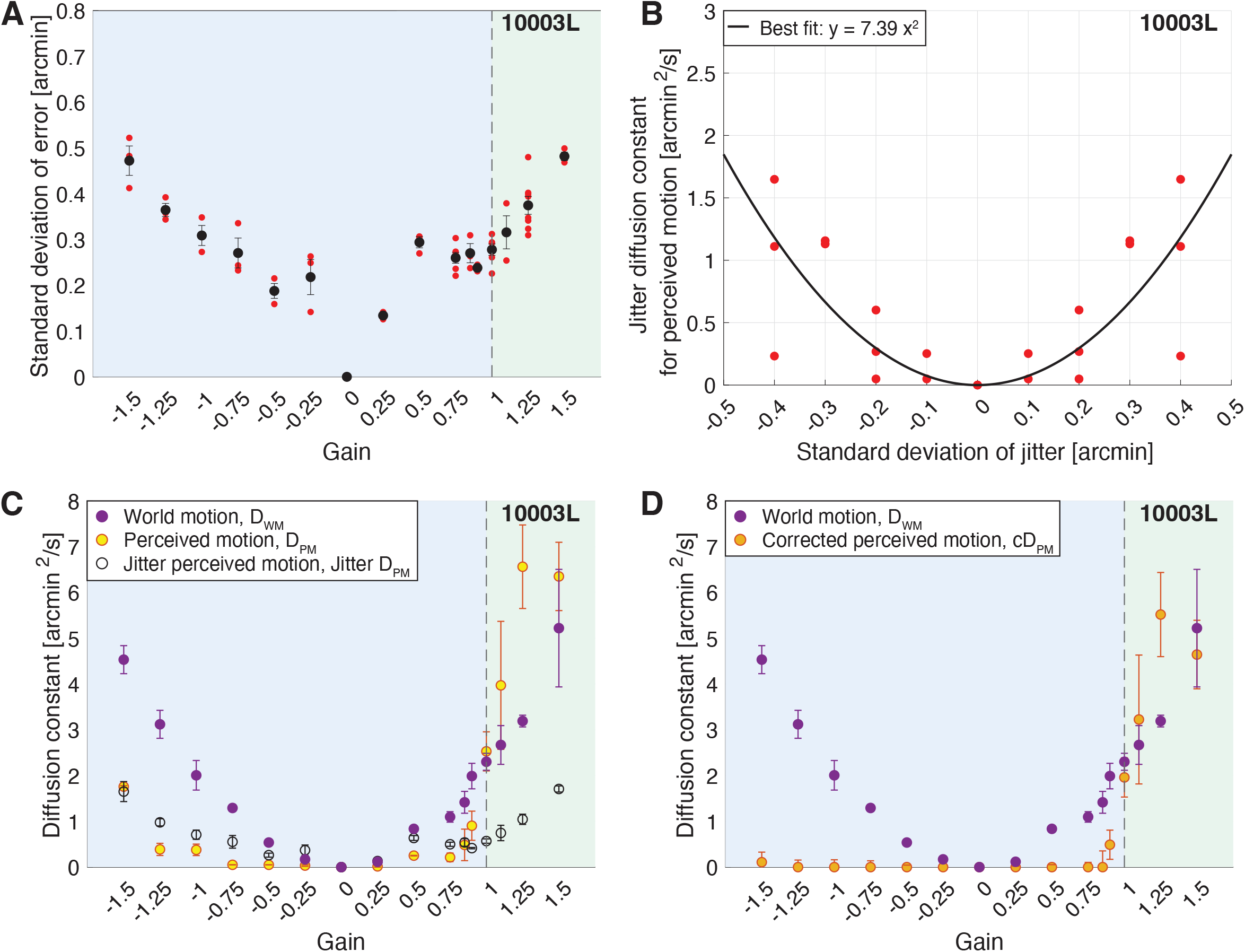
Results from subject 10003L. The blue region of graphs (A), (C) and (D) indicates the Gains where the stimulus’ motion across the retina is in a direction that is directly opposite to eye motion. The green region indicates the Gains where the stimulus’ motion across the retina is in the same direction as eye motion. (A) The magnitude of stimulus delivery error for each Gain. For each trial, we computed the the distances in x and y between the ideal retina-contingent stimulus motion trace and the measured retina-contingent stimulus motion trace. The red points indicate the standard deviation of error from one trial – this is the standard deviation across the distances in x and y from all frames from all videos within a single trial. The black points indicate the average standard deviation of error across all trials, with SE of the mean bars. (B) Jitter diffusion constants for perceived motion, *Jitter D*_*PM*_, plotted as a function of the standard deviation which was used to generate the jitter stimulus. Each red point represents one trial and was reflected across the y-axis. The black curve is a best-fit quadradic function which is anchored at 0 arcmin^2^/s. (C-D) Diffusion constants plotted as a function of the sixteen Gains. Purple points indicate average diffusion constants for world motion, *D*_*WM*_, with SE of the mean bars. (C) Yellow points indicate average diffusion constants for perceived motion, *D*_*PM*_, with SE of the mean bars. Black-hollow points indicate average *Jitter D*_*PM*_ with SE of the mean bars, given the standard deviation of error in (A). (D) Orange points indicate average corrected diffusion constants for perceived motion, *cD*_*PM*_, with SE of the mean bars. *cD*_*PM*_ = *D*_*PM*_ − *Jitter D*_*PM*_.

Overall, the AOSLO real-time tracking system demonstrates subarcminute tracking accuracy. It is noted here that tracking can never be perfect because of the ~2-ms latency between the last measurement of eye position and the delivery of the stimulus (Arathorn et al., 2007).

### Quantifying the Perceived Motion of the Retina-Contingent Stimuli

After the experiment, we recovered the random walk stimulus traces at the setting of a perceptual match and computed a diffusion constant across these traces. The diffusion constant of the random walk stimulus represents the magnitude of motion that the subject perceived when they viewed the retina-contingent stimulus. We will refer to this value as the diffusion constant for perceived motion, or *D*_*PM*_. An example from one subject is shown in Fig. 2C (yellow points) and for all subjects in Fig. A1.2 in Appendix 1.

#### Accounting for Motion Perception Due to Error

Next, we were motivated to determine to what degree stimulus delivery errors, which were computed in Section **Quantifying the Stimulus Delivery Error**, contributed to the subjects’ motion judgments during the experiments. These errors manifest as approximately random, non-cumulative steps, or “jitter”, which are superimposed onto the retina-contingent stimulus’ motion. If the visual system indeed fully suppresses the perception of motion that is in a direction consistent with the direction of retinal slip, then it is possible that under these conditions, the motion that the subjects matched was solely influenced by the jitter due to the tracking errors.

To measure the perceived motion of jitter, we executed the same method-of-adjustment procedure, except that we replaced the retina-contingent stimulus with a stimulus that moved independently of eye motion. This stimulus jittered in place with step sizes in x and y that were determined by a standard deviation. We tested the following standard deviations: 0, 0.1, 0.2, 0.3, and 0.4 arcmin, refer to Section **Generating Random Walks and Random Jitter** for more details on the generation of this stimulus. Note that because subject 20237R’s average standard deviations of error were below 0.4 arcmin, we did not test this subject with a 0.4 arcmin standard deviation of jitter, shown in Fig. A1.1 in Appendix 1. Subjects were instructed to adjust the magnitude of motion of the random walk stimulus until it matched the perceived motion of the jitter stimulus. We computed a diffusion constant of the random walk stimulus at the setting of a perceptual match which represents the subjects’ perceived motion of the jitter stimulus. This value will be referred to as the Jitter diffusion constant for perceived motion, or *Jitter D*_*PM*_. The subjects performed three trials for each standard deviation condition. Fig. 2B plots the *Jitter D*_*PM*_ as a function of the standard deviations for one subject. We reflected the results across the y-axis and fit a quadratic function across the data points. Because perceived motion cannot be negative, we anchored the function to 0 arcmin^2^/s using the MATLAB function polyfitZero (Mikofski, 2025). Results for all subjects are reported in Fig. A1.1 in Appendix 1.

Using methods described in Section **Quantifying the Stimulus Delivery Error**, for each subject, we determined the magnitude of stimulus delivery error for each Gain (Fig. 2A). Using the subjects’ perceptual responses of the jitter stimulus (black curve in Fig. 2B), we estimated the perceived motion of this stimulus delivery error (black points in Fig. 2A). For example, for 10003L, the average standard deviation of stimulus delivery error during presentations of Gain −1.5 stimuli was ~0.47 arcmin (Fig. 2A) which corresponded to an approximate perceived motion of 1.65 arcmin^2^/s (Fig. 2B).

The following were computed for each subject. For each Gain, we averaged the diffusion constants for perceived motion of the retina-contingent stimulus (average *D*_*PM*_, Fig. 2C yellow points) and we averaged the perceived motion of error (*Jitter D*_*PM*_, Fig. 2C, black-hollow points). We subtracted average *D*_*PM*_ − average *Jitter D*_*PM*_, and the result will be referred to as the “corrected diffusion constant for perceived motion”, or *cD*_*PM*_. Negative *cD*_*PM*_ values were set equal to 0 because motion perception cannot be negative. Fig. 2D shows the average *cD*_*PM*_ for one subject and Fig. 3A and Fig. A1.2 in Appendix 1 show the average *cD*_*PM*_ for all subjects.

**Figure 3:**
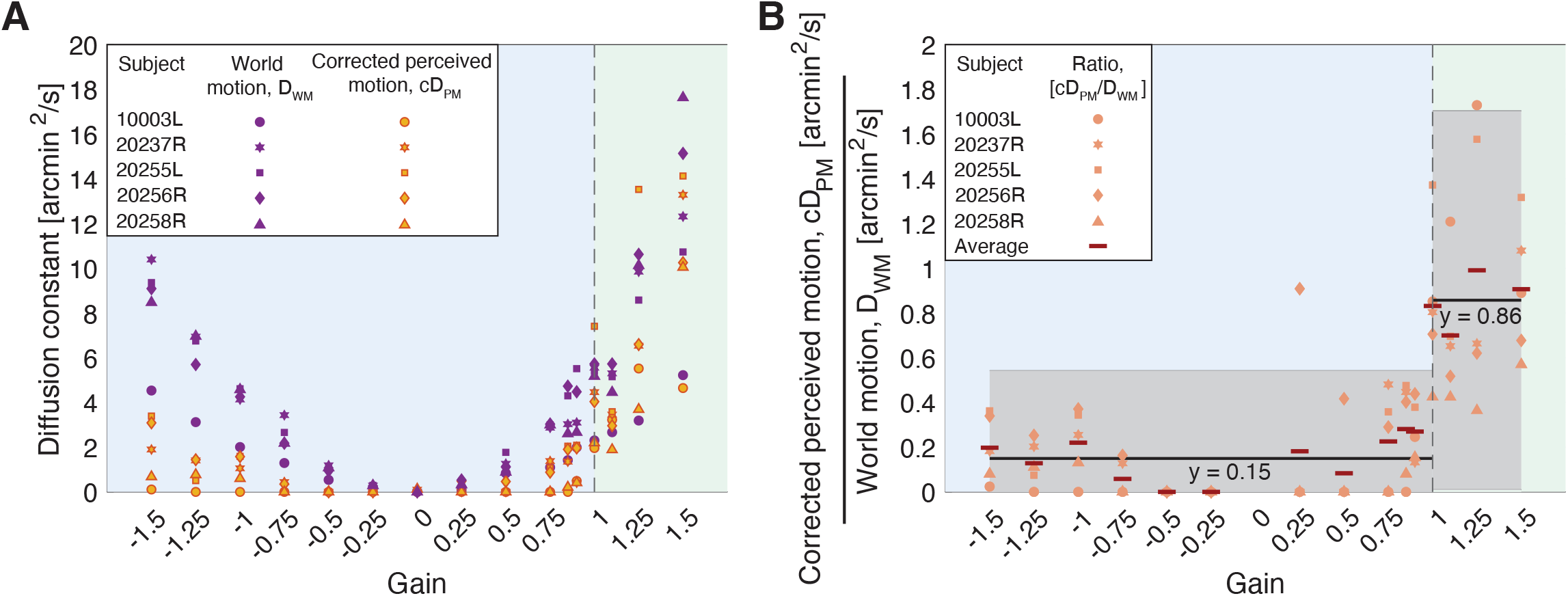
Experiment 1: Perceived motion and world motion of the retina-contingent stimuli. The blue regions indicate the Gains where the stimulus’ motion across the retina is in a direction that is directly opposite to eye motion. The green regions indicate the Gains where the stimulus’ motion across the retina is in the same direction as eye motion. (A) Diffusion constants from five subjects as a function of the Gains. Orange points indicate average corrected diffusion constants for perceived motion, *cD*_*PM*_. Purple points indicate average diffusion constants for world motion, *D*_*WM*_. Each point represents one subject and is the average across all matches. (B) Average *cD*_*PM*_ */D*_*WM*_ from five subjects as a function of the Gains. Each point represents one subject and is the average ratio across all matches. The red bars indicate the average ratio across all subjects. Note that ratios for Gain 0 were removed. We performed independent, single-parameter linear fits to the ratios of Gains *<* +1 (*y* = 0.15) and to the ratios of Gain ≥ +1 (*y* = 0.86). The gray intervals indicate the 95% confidence bounds of the fits.

### Quantifying the World Motion of the Retina-Contingent Stimuli

We collected the retina-contingent stimulus motion traces from every video within a trial, excluding videos that were removed after the filtering steps in Section **Quality Control**. We computed the diffusion constant across all traces within each trial and this value will be referred to as the diffusion constant for world motion of the retina-contingent stimulus, or *D*_*WM*_. Fig. 2 C and D show the average *D*_*WM*_ for one subject and Fig. 3A and Fig. A1.2 in Appendix 1 show the average *D*_*WM*_ for all subjects. Observe that while the instantaneous speed of the stimulus motion increases linearly with the magnitude of the Gain, the associated diffusion constants increase with the square of the Gain.

### Ratio Analysis

For each subject we performed the following: For each Gain, we computed the ratio of the [average corrected diffusion constant for perceived motion] to the [average diffusion constant for world motion], or average *cD*_*PM*_ */D*_*WM*_. Because world-fixed stimuli have *D*_*WM*_ = 0 arcmin^2^/s, we removed the ratios for Gain 0. Fig. 3B shows the average *cD*_*PM*_ */D*_*WM*_ for each subject, and Fig. A1.3 in Appendix 1 reports the individual trial distributions of *cD*_*PM*_ */D*_*WM*_ for all subjects. We expect the ratio to be one if subjects perceived the motion veridically. Ratios less than one indicate that subjects underestimated the motion and ratios greater than one indicate that subjects overestimated the motion.

### Model Comparison Metric

We performed multiple function fits to the ratios in Fig. 3B to assess the sharpness of the discontinuity in motion perception during drift, where *x* represents the Gain and *y*_*i*_ represents the average *cD*_*PM*_ */D*_*WM*_. If the discontinuity is sharp, then the data would be best fit with two independent functions on either side of a breakpoint. To evaluate the different fits, we computed the least squares (corrected) Akaike information criterion or *AIC*_*c*_. This model comparison metric evaluates goodness-of-fit while penalizing models with increasing number of parameters. As recommended by Burnham and Anderson (2002), the AIC is added to a penalty term to reduce small-sample bias. After computing the *AIC*_*c*_ for each model, we computed evidence ratios to assess the strength of evidence for each model, using methods described in Burnham and Anderson (2002). Higher evidence ratios indicate stronger evidence for the “best” model. Lower evidence ratios indicate weaker support for the “best” model and suggests more uncertainty for model selection (Burnham & Anderson, 2002). An extended description of the models and the equations for the *AIC*_*c*_ and evidence ratio are reported in Appendix 1.

### Statistics

For Experiment 2, we performed a 2-factor, repeated-measures ANOVA of the diffusion constant for eye motion (*D*_*EM*_), *α* for eye motion (*α*_*EM*_), and corrected diffusion constant for perceived motion (*cD*_*PM*_). There were 3 Gains (−1.5, 0, and +1.5) and three background conditions (no-white-circle, 2°-diameter white circle, and 9°-diameter white circle conditions), which were within-subject factors. If the repeated-measures ANOVA found an effect, indicated by *P <* 0.05, we performed a post hoc Tukey-Kramer test to measure the significance of the pairwise comparisons.

## Results

### Experiment 1

Fig. 3A plots the average diffusion constants as a function of the sixteen Gains for the five subjects. The orange points indicate average corrected diffusion constants for perceived motion, *cD*_*PM*_, of the retina-contingent stimuli. The purple points indicate average diffusion constants for world motion, *D*_*WM*_, of the retina-contingent stimuli. Each point represents one subject and is the average across at least three trials.

Fig. 3B plots the average *cD*_*PM*_ */D*_*WM*_ as a function of the sixteen Gains for the five subjects. A *cD*_*PM*_ */D*_*WM*_ equal to one indicates that the subject perceived the stimulus’ actual motion in the world, a *cD*_*PM*_ */D*_*WM*_ lower than one indicates that the subject underestimated the world motion, and a *cD*_*PM*_ */D*_*WM*_ greater than one indicates that the subject overestimated the world motion.

#### A Sharp Discontinuity in Motion Perception During Drift

We performed multiple fits to the ratios in Fig. 3B to assess the sharpness of the discontinuity in motion perception during drift. We tested linear fits on either side of a breakpoint, the breakpoints ranged from Gain −1 to +1.1. We tested single-parameter linear fits as well as two-parameter linear fits in slope-intercept form. We also tested a single fit across all ratios: a quadradic, a cubic, and a quartic function. Examples are shown in Fig. 3B and Fig. A1.4 of Appendix 1.

The corrected Akaike information criterion (*AIC*_*c*_) values were lowest (“best”) for the single-parameter linear fits with a breakpoint at Gain +1. The two-parameter linear fits with a breakpoint at Gain +1 had the second lowest *AIC*_*c*_ value, although the evidence ratio is ~3.89 in relation to the single-parameter linear fit model, which suggests that there is little evidence in favor of this model. The *AIC*_*c*_ values for all other models were more positive with evidence ratios greater than 150, meaning that evidence is reasonably strong against these models. Thus, given the models tested and the dataset, the single-parameter linear fits with a breakpoint at Gain +1 best fit the data suggesting that there is reasonably strong evidence supporting a sharp discontinuity in motion perception during drift where direction with respect to eye motion is the critical parameter in governing how motion is perceived. Stimuli moving opposite to eye motion (Gains *<* 1) are underestimated (~0.15x), while stimuli moving in the same direction as eye motion (Gains ≥ 1) are perceived as moving, and the motion perceived is similar to the stimulus’ actual motion in the world (~0.86x). Model comparison metrics for all models are reported in Table A1.1 in Appendix 1. We additionally performed the same calculations using the original, uncorrected diffusion constants for perceived motion (*D*_*PM*_) in place of the *cD*_*PM*_ of the ratio, and we found the same trends, which are reported in Table A1.2 in Appendix 1.

### Experiment 2

Fig. 4 plots the average diffusion constants for each of the three Gains tested (Gains −1.5, 0 and +1.5) under three background conditions (no-white-circle, 2°-diameter white circle, and 9°-diameter white circle conditions). Because there were only three Gains, we plotted the average corrected diffusion constants for perceived motion (*cD*_*PM*_) as a function of the average diffusion constants for world motion (*D*_*WM*_). If the subjects perceived the actual motion of the stimulus in the world, then the data points would lie along the 1:1 line. The small symbols represent each subject’s average perceptual match for Gain −1.5 stimuli (blue open symbols), Gain 0 stimuli (yellow filled symbols), and Gain +1.5 stimuli (green filled symbols). The large stars are the group averages with SE of the mean bars. The red arrows show the extent to which the eye motion, and consequent retina contingent stimulus’ world motion (*α*_*WM*_), deviated from Brownian. Arrows pointing Right indicate persistence (*α*_*WM*_ *>* 1), arrows pointing Left indicate antipersistence (*α*_*WM*_ *<* 1), and no arrow means that the motion was Brownian (*α*_*WM*_ = 1 +*/*− 0.02). We did not show red arrows for the Gain 0 stimuli because motion statistics do not apply to world-fixed stimuli. As discussed in D’Angelo et al. (2024), the presence of persistent eye motion generally gives rise to relatively higher values of the *D*_*WM*_ and relatively lower values of the diffusion constant of the random walk at the perceived match (*D*_*PM*_). As a result, subjects with higher degrees of persistence tend to plot below the 1:1 line. Note that subject 20256R was not tested under the no-white-circle condition. Fig. A1.5 of Appendix 1 reports individual trial distributions of the matches for all subjects, and Fig. A1.6 of Appendix 1 reports the standard deviation of stimulus delivery error distributions for all subjects.

**Figure 4:**
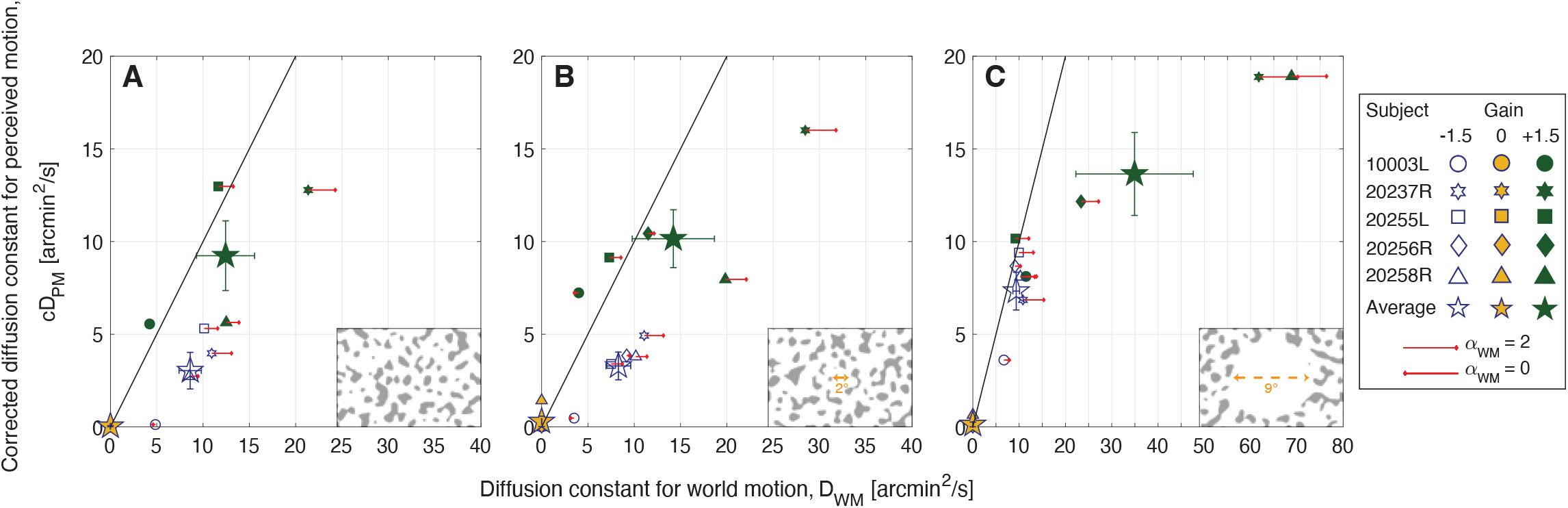
Experiment 2: Average corrected diffusion constants for perceived motion (*cD*_*PM*_) versus diffusion constants for world motion (*D*_*WM*_) from five subjects. Experiments were tested under three background conditions: The patterns extended over the full 17° field of view (A), or surrounded a 2°-diameter white circular opening (B), or surrounded a 9°-diameter white circular opening (C). Background conditions are indicated by labels on the Lower Right corner of each graph. Note that subject 20256R was not tested under the no-white-circle condition. The small symbols represent each subject’s average perceptual match for Gain −1.5 stimuli (blue open symbols), Gain 0 stimuli (yellow filled symbols), and Gain +1.5 stimuli (green filled symbols). The large stars are the group averages with SE of the mean bars. The red arrows show the extent to which the eye motion, and consequent retina contingent stimulus’ world motion (*α*_*WM*_), deviated from Brownian. Arrows pointing Right indicate persistence (*α*_*WM*_ *>* 1), arrows pointing Left indicate antipersistence (*α*_*WM*_ *<* 1), and no arrow means that the motion was Brownian (*α*_*WM*_ = 1 +*/*− 0.02). Longer arrows correspond to higher deviations from Brownian motion. The arrow length in the legend indicates pure persistence (*α*_*WM*_ = 2, straight line trajectory at constant velocity) if pointing Right or pure antipersistence (*α*_*WM*_ = 0, oscillatory motion) if pointing Left.

#### The Illusion of Relative Stability Depends on the Extent of Retinal Image Background Content

The illusion of relative stability persisted even when the retinal image background content extended into – and even overlapped with – the retina-contingent stimulus. There was no statistically significant difference in perceived motion of Gain −1.5 stimuli between the no-white-circle and 2°-diameter white circle conditions (Fig. 4 A and B). For both backgrounds, Gain +1.5 stimuli were perceived to have more motion than Gain −1.5 stimuli (*P* = 0.036 for no-white-circle condition and *P* = 0.037 for 2°-diameter white circle condition, post hoc Tukey–Kramer). This is surprising considering that, under the no-white-circle condition, the stimulus in many instances would have crossed from dark to light background regions while the matches were being made.

On the other hand, subjects perceived significantly higher magnitudes of motion of Gain −1.5 stimuli when the retinal image background content was further displaced (the 9°-diameter white circle condition shown in Fig. 4C) from the retina-contingent stimulus (*P* = 0.01 compared to the no-white-circle condition and *P* = 0.04 compared to the 2°-diameter white circle condition, post hoc Tukey–Kramer). By spatially offsetting the background content ~4° from the stimuli, subjects perceived significantly higher magnitudes of motion of Gain −1.5 stimuli and perceptually underestimated the motion of Gain +1.5 stimuli (Fig. 4C), results that lean towards those tested in a Ganzfeld reported in a prior paper (D’Angelo et al., 2024).

For Gain +1.5 stimuli, perceived motions were not significantly different between any background conditions (Fig. 4 A−C).

As an additional check, we performed the repeated-measures ANOVA and post hoc Tukey-Kramer tests using the original, uncorrected diffusion constants for perceived motion (*D*_*PM*_) for all Gains and background conditions from Experiment 2 and these trends held.

### Additional Analyses: Effect of Gain on Eye Motion Statistics

Fig. 5 plots the average diffusion constants for eye motion (*D*_*EM*_) and average *α* for eye motion (*α*_*EM*_) for all subjects as a function of the Gains tested in Experiments 1 and 2. Within the violins (Bechtold, 2016), the central white circles represent the median and the dark gray bars represent the interquartile range. The surrounding intervals are the density traces (Hintze & Nelson, 1998).

**Figure 5:**
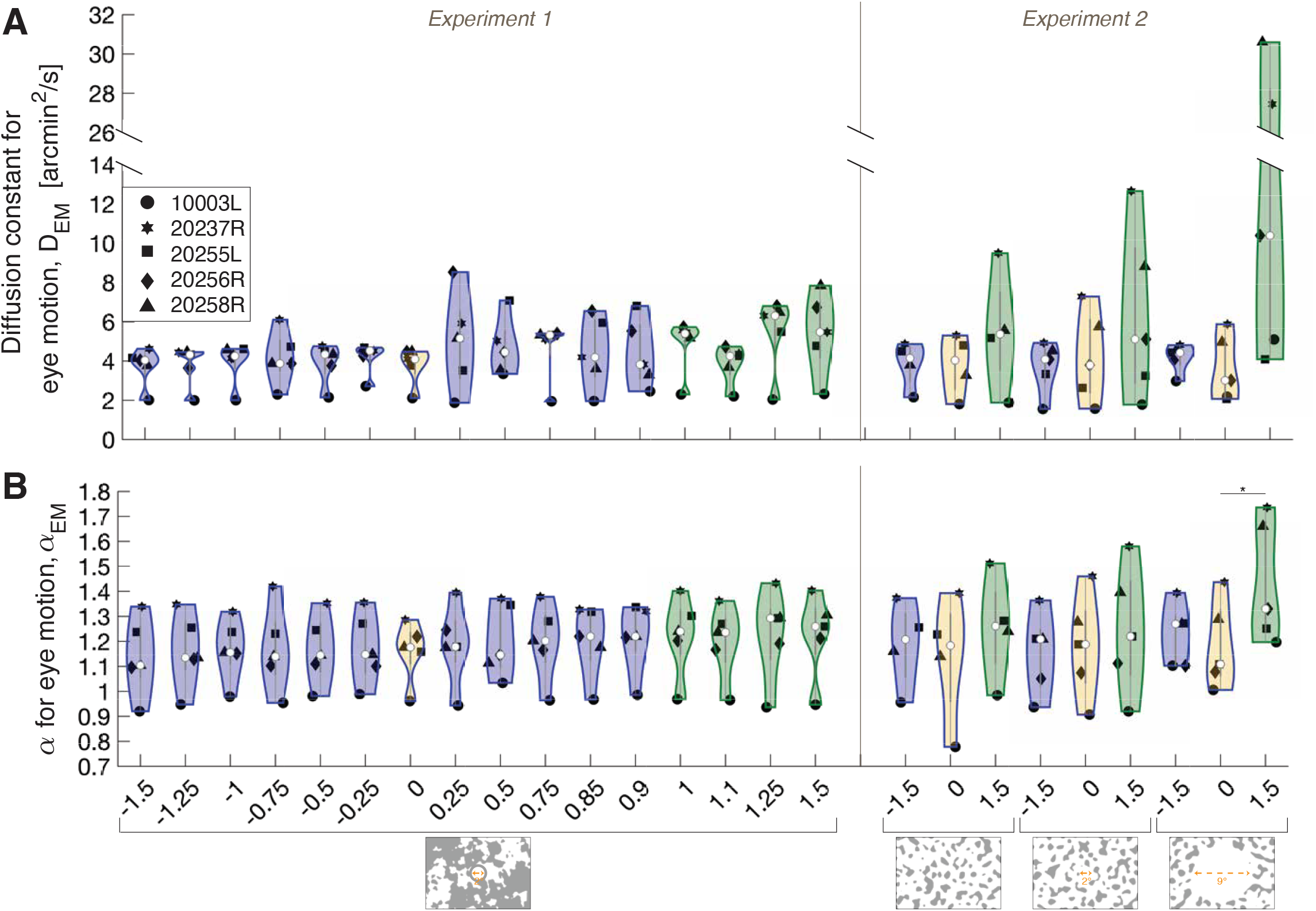
Violin plots showing the distribution of the (a) diffusion constant for eye motion, *D*_*EM*_ and (b) *α* for eye motion, *α*_*EM*_ as a function of the Gains from Experiments 1 and 2. Background conditions are indicated by labels at the bottom of the graph. The violins outlined in blue (including the yellow-filled violin which labels the world-fixed stimulus) indicate the Gains where the stimulus’ motion across the retina is in a direction that is directly opposite to eye motion. The violins outlined in green indicate the Gains where the stimulus’ motion across the retina is in the same direction as eye motion. Within each violin, the central white circles represent the median and the dark gray bars represent the interquartile range. The surrounding intervals are the density traces. Each black point represents one subject and is the (a) average *D*_*EM*_ and (b) average *α*_*EM*_ across their trials for each Gain. The asterisk indicates statistical significance (*P <* 0.05) from a post hoc Tukey-Kramer test following a two-factor repeated-measures ANOVA.

#### Experiment 1

Undergoing the same eye motion, increasing the magnitude of the Gain gives rise to higher magnitudes of the stimulus’ world motion. Therefore, we computed the ratio [average corrected diffusion constant for perceived motion]/[average diffusion constant for world motion], or *cD*_*PM*_ */D*_*WM*_, to account for the world motion. We were motivated to ensure that Gain conditions did not impact the eye motion statistics, in particular the diffusion constant for eye motion, *D*_*EM*_ (Fig. 5A, Experiment 1) and *α* for eye motion, *α*_*EM*_ (Fig. 5B, Experiment 1). We found no significant difference between *D*_*EM*_ for Gains *<* +1 compared to *D*_*EM*_ for Gains ≥ +1 (*P* = 0.092, Paired t-test). We found no significant difference between *α*_*EM*_ for Gains *<* +1 compared to *α*_*EM*_ for Gains ≥ +1 *α*_*EM*_ (*P* =0.12, Paired t-test).

#### Experiment 2

A repeated measures ANOVA revealed no significant effect of Gain or background condition on *D*_*EM*_ (*P* = 0.11 and *P* = 0.085, respectively), however the interaction of Gain and Background had a significant effect on *D*_*EM*_ (*P* = 0.034). A post hoc Tukey Kramer showed no significant differences between pairwise comparisons of *D*_*EM*_ (Fig. 5A, Experiment 2). This is likely due to small effect sizes between conditions which are not robust to the post hoc correction, given the current sample size (n = 5 subjects).

Additionally, a repeated measures ANOVA revealed a significant effect of Gain on *α*_*EM*_ (*P* = 0.030) but no significant effect of background condition on *α*_*EM*_ (*P* = 0.094). The interaction of Gain and Background had a significant effect on *α*_*EM*_ (*P* = 0.018). Under the 9°-diameter white circle condition, the *α*_*EM*_ of Gain +1.5 stimuli were significantly higher than those of Gain 0 stimuli (Fig. 5B, Experiment 2).

Thus, the perceived motion (*cD*_*PM*_) cannot be explained by changes in eye motion statistics, *D*_*EM*_ and *α*_*EM*_, across Gain conditions. Offsetting the background content and turning off the fixation cross (9°-diameter white circle condition) can explain the higher *D*_*EM*_ and *α*_*EM*_ (Fig. 5 A and B, Experiment 2), which is consistent with a previous report (D’Angelo et al., 2024).

## Discussion

We demonstrate that perception of moving objects during fixational drift is discontinuous, governed by the direction those objects move with respect to eye motion. Our results build on previous studies (Arathorn et al., 2013; D’Angelo et al., 2024) and show that the direction that objects move with respect to eye motion, regardless of magnitude, is a critical parameter that governs an exception to our ability to detect relative motion.

### Two Paradoxes in Motion Perception During Drift

The human visual system relies on paradoxical methods to veridically perceive stable and moving objects.

Paradox 1: To perceive world-fixed objects in high resolution, the eye attempts to stabilize images-of-interest onto the fovea (Epelboim & Kowler, 1993; Clark et al., 2022); yet, paradoxically, if the eye achieves perfect stability the image will quickly fade from view (Ditchburn & Ginsborg, 1952; Riggs & Ratliff, 1952). The eye therefore is in constant motion, and the human visual system evolved to discount the image motion due to eye motion, and even leverages its motion to improve spatial vision (Ratnam, Domdei, Harmening, & Roorda, 2017; Rucci, Iovin, Poletti, & Santini, 2007).

Paradox 2: The visual system relies on world-fixed frames of reference to properly detect moving objects in the world (Legge & Campbell, 1981). However, the same frames of reference serve as retinal image background content to perceptually stabilize images that move in a direction consistent with the direction of retinal slip, regardless of magnitude, giving rise to the illusion of relative stability (D’Angelo et al., 2024).

### Where in the Visual Pathway Does the Illusion Take Place?

As discussed in D’Angelo et al. (2024), the mechanisms underlying the illusion of relative stability at the earliest stages may rely on signals from the directionally sensitive retinal ganglion cells (DSRGCs) whose existence in primate have been recently reported (A. Y. Wang et al., 2023). As for the downstream processing of these signals, a functional circuit-level hypothesis is being developed to explain how the behavior reported here and in previous papers might arise.

While the exact mechanisms remain unknown, the current study adds three important factors to consider: First, the illusion persists for Gains between 0 and +1, where the retinal image slip is consistent with, but less than, the eye’s motion (Fig. 3). Second, the illusion is so profound that it persists even when the retinal image background content overlaps entirely with the retina-contingent stimuli (Fig. 4A). Third, if the retinal image background content is too far from the retina-contingent stimuli, then the effect diminishes (Fig. 4C), suggesting that the effect might not be a true discontinuity, but rather a graded response. To further elucidate possible mechanisms, the dependence on proximity to retinal image background content, and also the dependence on spatial frequency of that content can be explored. These remain parameters for future study.

### Evaluating the Sharpness of the Discontinuity

As mentioned above, the illusion of relative stability may not be a true discontinuity, but rather a graded response. We will review several reasons: First, if the retinal image background content is too far from the retina-contingent stimuli, then the effect diminishes (Fig. 4C) suggesting that the magnitude of this effect is driven by proximity to retinal image background content. Second, if it were a true discontinuity, the perceived motion (diffusion constant for perceived motion, *D*_*PM*_) for Gains *<* +1 would be close to zero, regardless of Gain magnitude. However, all subjects showed a slightly increasing perception of motion (higher *D*_*PM*_) as the Gains approached one or increased in retinal slip (Fig. A1.2 in Appendix 1). Third, in our experiments, all motion was restricted in two dimensions to be always aligned with the eye motion, either in the same direction or opposite; orientations of motion in other directions not aligned with eye motion also show a more gradual transition (Arathorn et al., 2013).

Further, even after accounting for stimulus delivery errors by computing the corrected diffusion constant for perceived motion, *cD*_*PM*_, the perceived motion of stimuli moving opposite to eye motion (Gains *<* +1) did not go down to zero for most subjects. All subjects perceived slightly increasing magnitudes of motion as the Gains approached one or increased in retinal slip (Fig. A1.2 in Appendix 1). This suggests that the motion the subjects saw was not solely due to stimulus delivery errors. And this suggests that higher magnitudes of stimulus motion, compared to the magnitude of eye motion, serve to diminish the illusion, supporting that this is a graded response. While the *D*_*PM*_ and *cD*_*PM*_ increased slightly with Gain magnitude for Gains *<* +1, it is important to note that the *D*_*PM*_ and *cD*_*PM*_ remained well below the magnitude of the stimulus’ motion in the world (diffusion constant for world motion, *D*_*WM*_).

### Comparison with Previous Work

In Arathorn et al. (2013) and D’Angelo et al. (2024), under background-present conditions, stimuli moving in the same direction as eye motion (Gains ≥ +1) appeared readily in motion. For Gains ≥ +1, we would expect that the motion the subjects perceived would be equal to the motion of the stimuli in the world. Arathorn et al. (2013) did not quantify the magnitude of eye motion or the retina contingent stimulus’ world motion so we cannot make any claims on whether the subjects veridically perceived the world motion of the retina-contingent stimuli. But D’Angelo et al. (2024) computed the diffusion constant and *α* of the eye motion (*D*_*EM*_ and *α*_*EM*_) and retina-contingent stimulus’ world motion (*D*_*WM*_ and *α*_*WM*_); the same method is used in this study, so we can compare the two studies as shown in Fig. A2.1A in Appendix 2. The results are generally the same but the differences in the experimental parameters and possible consequences of those are discussed in Appendix 2.

In D’Angelo et al. (2024), we report results from experiments under a Ganzfeld condition. In the current study, as a control, we did the same method-of-adjustment experiments under a Ganzfeld as well, and we found the same trends: that motion perceptions reversed in the Ganzfeld. Similar to D’Angelo et al. (2024), *α*_*EM*_ was significantly higher under the Ganzfeld (*P <* 0.001, Paired t-test). Results from those experiments are included in Fig. A1.7 of Appendix 1.

### Causal Inference in Perception

The discontinuity that leads to the illusion of relative stability may arise from a causal inference process where, before estimating the actual motion of the stimuli, the visual system must decide if a given object is part of the world-fixed background or a moving object. Therefore, the illusion is in line with the notion of a visual system that has evolved to discount images moving with amplified retinal slip, as they are more likely to be part of the world-fixed stimuli. Given that it is highly unlikely that any object would ever move with amplified retinal slip for an appreciable time relative to other retinal image background content in the visual scene, it makes sense that such motion should never contaminate our perception. A causal inference framework could be used to formalize and explain this process. Causal inference has been used to explain perceptual processing in a growing number of studies (Shams & Beierholm, 2010; Dokka, Park, Jansen, DeAngelis, & Angelaki, 2019; Acerbi, Dokka, Angelaki, & Ma, 2018). Future work could uncover a causal inference model in motion perception during drift.

## Conclusions

We showed that during fixational drift, the direction an object moves with respect to eye motion is a crucial parameter in giving rise to perceptions of stability and motion. Images moving opposite to eye motion, in a direction consistent with retinal slip, were perceived as relatively stable while the motion of images moving in the same direction was readily detected. This held even for stimuli that were slipping opposite to eye motion but with a smaller magnitude across the retina. We determined that this misperception persists even if the image moves on top of or close to retinal image background content – which is surprising and paradoxical given how this same content is used as a frame of reference to detect relative motion of images moving in any other direction with respect to eye motion.

## Acknowledgments

The authors thank Dr. Benjamin M. Chin for his helpful discussions. This research was supported by funding from National Institutes of Health Grants R01EY023591 (A.R.) and T32 EY007043 (J.C.D.), the Center for Innovation in Vision and Optics (civo.berkeley.edu) (J.C.D.), and a grant from the Minnie and Roseanna Turner Fund for Impaired Vision Research (J.C.D.).

Commercial relationships: P.T. and A.R. are co-inventors on US Patent #10130253, assigned to the University of California.

## Appendix 1

### Linear and Nonlinear Fits

For the data plotted on Figure 3B, we tested multiple breakpoints ranging from Gain −1 to +1.1, and performed an independent linear fit to each side of the breakpoint, where *x* represents the Gain and *y*_*i*_ represents the [average corrected diffusion constant for perceived motion]/[average diffusion constant for world motion], or *cD*_*PM*_ */D*_*WM*_.

We tested independent, single-parameter linear fits on either side of the break point.

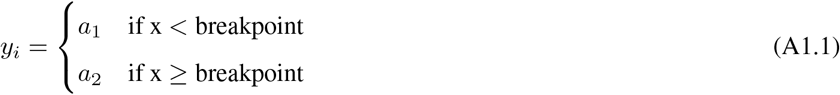

We also tested independent, two-parameter linear fits in slope-intercept form on either side of the breakpoint.

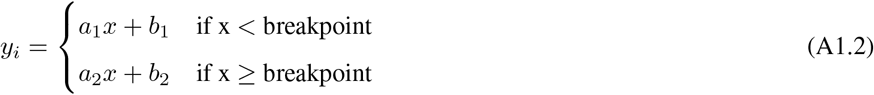

If the discontinuity is weak, then the data would be best fit with a single function across all data points. We tested a quadradic, a cubic, and quartic function.

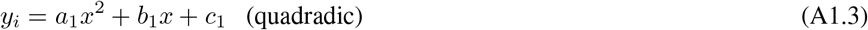

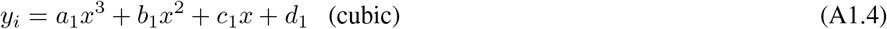

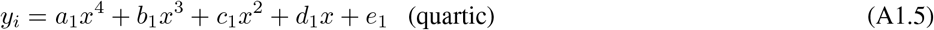

The fit across all Gains, *y*_*i*_, is expressed in Equation A1.1−A1.5 and will be referred to as a “model”.

### Akaike Information Criterion

The Akaike information criterion (AIC) formula is expressed in Equation A1.6, where n is the sample size and 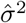 is the average sum of the squared residuals or mean squared error (Equation A1.7). K is the number of regression parameters, including the intercept and variance (Burnham & Anderson, 2002). As recommended by Burnham and Anderson (2002), the AIC is added to a penalty term to reduce small-sample bias, shown in Equation A1.8.

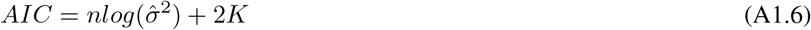

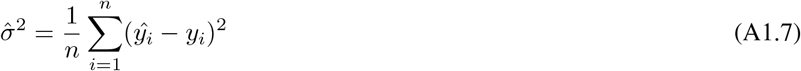

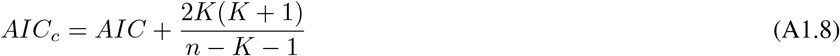

### AIC Interpretation: Computing Evidence Ratios

Lower *AIC*_*c*_ values are considered “better”, given the models tested and the dataset. To estimate if the differences in *AIC*_*c*_ values are meaningful, we performed the following as suggested by Burnham and Anderson (2002).

1. Subtracted the lowest 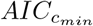 from each models’ 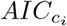.

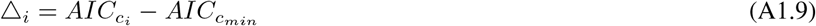
2. Computed the likelihood of each model *g*_*i*_ given the same dataset x.

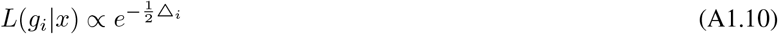
3. Computed “Akaike weights” *w*_*i*_ to determine the probability of each model *g*_*i*_ given R models tested.

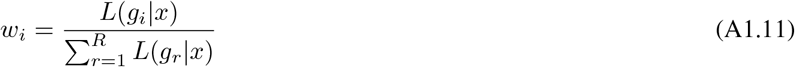
4. Computed “evidence ratios,” comparing each model’s Akaike weight *w*_*i*_ to the “best” model’s Akaike weight *w*_1_.

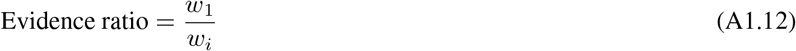

The *w*_*i*_ with the highest value is called *w*_1_. This means that there is strongest evidence that this is the “best” model, given R models tested. The evidence ratio is *w*_1_/*w*_*i*_, and therefore higher evidence ratios indicate stronger evidence for the “best” model. Lower evidence ratios indicate weaker support for the “best” model and suggests more uncertainty for model selection (Burnham & Anderson, 2002).

### Control Experiment: Perceived Motion under Background-Absent (“Ganzfeld”) Conditions

As a control, we performed the same method-of-adjustment as in this current study, except that the stimuli were presented under background-absent (“Ganzfeld”) conditions. Using methods described in D’Angelo et al. (2024), we set up a white paper with an aperture in front of the display permitting only the AOSLO and projector beams to enter the eye. We used the same aperture size as the one used in D’Angelo et al. (2024). LEDs were positioned between the subject and the paper to illuminate the paper. Due to its close proximity to the eye, the natural blur of the aperture in the paper rendered the transition between the display and luminance-matched paper invisible. Subjects performed the same matching procedure – fixating on the 680-nm cross and then attending to two circular stimuli. Subjects matched the motion of the left (random walk) stimulus to that of the right (retina-contingent) stimulus. To create a full Ganzfeld effect, we turned off the fixation cross during frames in which the two circular stimuli were presented. Without a fixation cross, eyes generally exhibit faster drift and larger corrective microsaccades (Cherici, Kuang, Poletti, & Rucci, 2012). To mitigate the effects of this increased retinal motion on the tracking accuracy, stimuli were presented for 750-ms. We measured the perceived motion of multiple retina-contingent stimuli with Gains ranging from −1.5 to +1.5 in increments of 0.25. A total of thirteen Gains were tested and the subjects performed three trials per Gain (except for 20237R and 20255L, who only performed two trials of Gain +0.75; 20256R who only performed two trials of Gain −0.75; and 20258R who only performed two trials of Gain −1.5. For all other Gains, these subjects performed 3 trials). Results from five subjects are shown Fig. A1.7.

**Figure A1.1:**
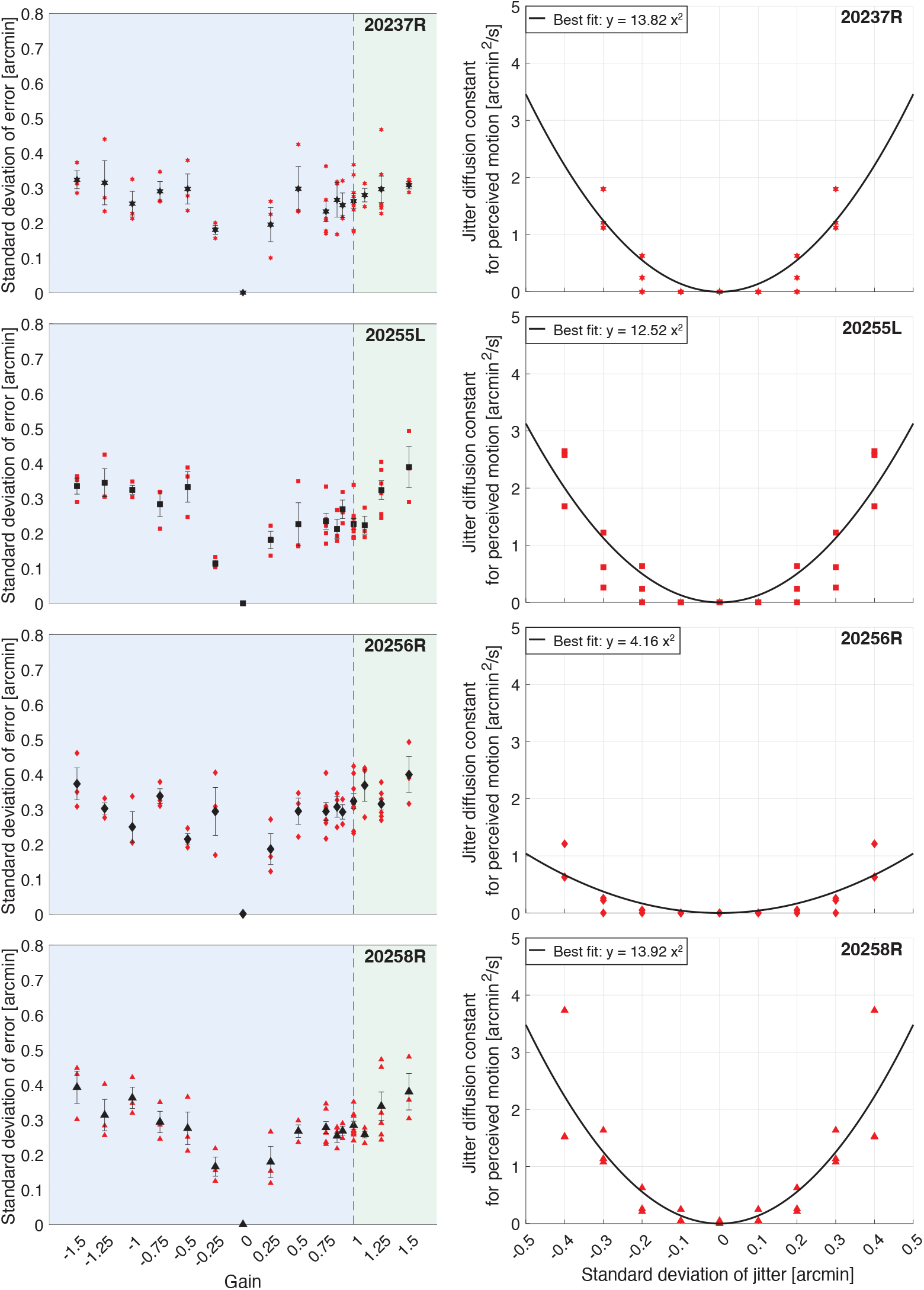
Experiment 1: Stimulus delivery error analysis from four subjects. The blue region of graphs indicates the Gains where the stimulus’ motion across the retina is in a direction that is directly opposite to eye motion. The green region indicates the Gains where the stimulus’ motion across the retina is in the same direction as eye motion. (**Left**) The magnitude of stimulus delivery error for each Gain. For each trial, we computed the the distances in x and y between the ideal retina-contingent stimulus motion trace and the measured retina-contingent stimulus motion trace. The red points indicate the standard deviation of error from one trial – this is the standard deviation across the distances in x and y from all frames from all videos within a single trial. The black points indicate the average standard deviation of error across all trials, with SE of the mean bars. (**Right**) Jitter diffusion constants for perceived motion, *Jitter D*_*PM*_, plotted as a function of the standard deviation which was used to generate the jitter stimulus. Each red point represents one trial and was reflected across the y-axis. The black curve is a best-fit quadradic function which is anchored at 0 arcmin^2^/s. Note that because subject 20237R’s average standard deviations of error were below 0.4 arcmin, we did not test this subject with a 0.4 arcmin standard deviation of jitter.

**Figure A1.2:**
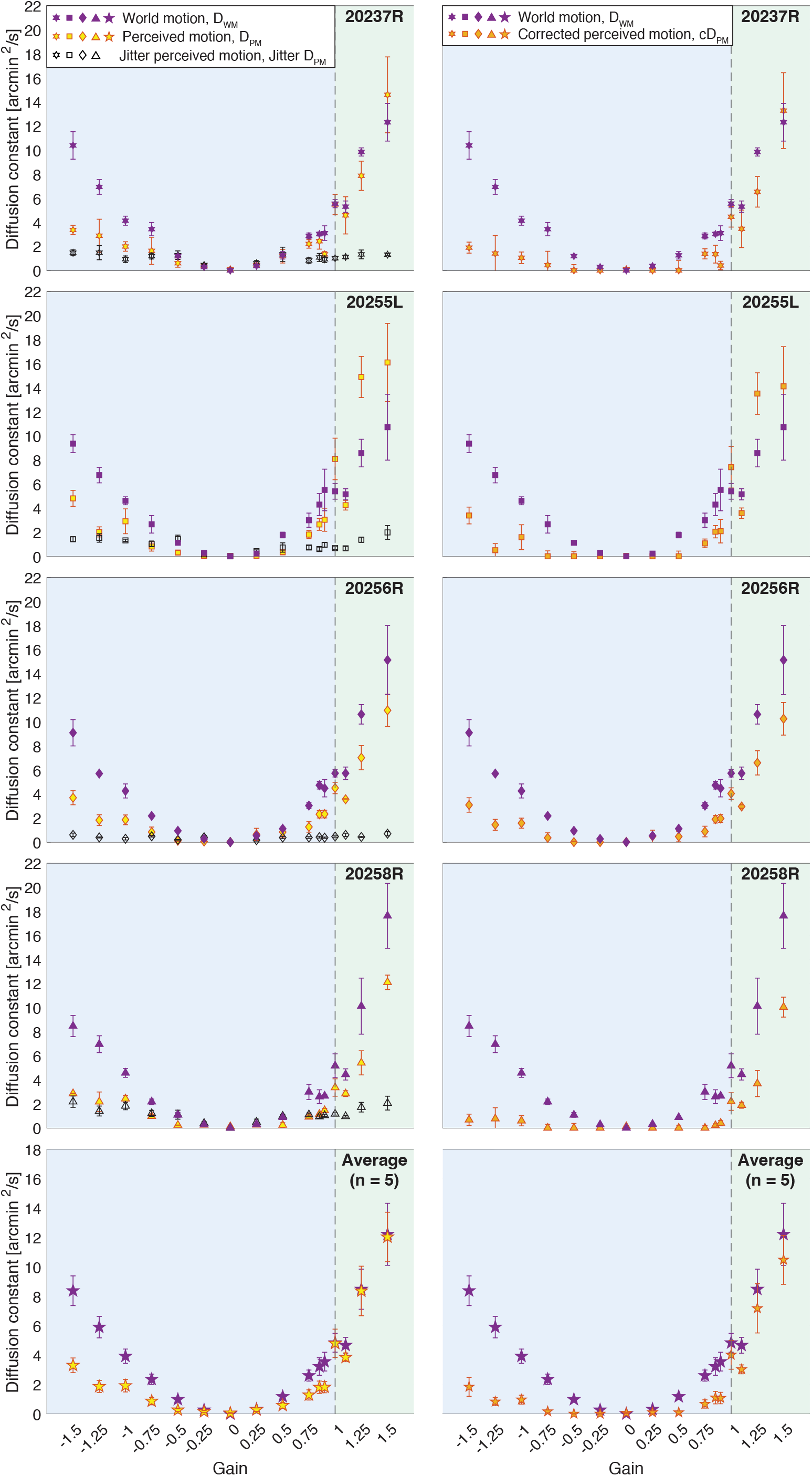
Experiment 1: Diffusion constants plotted as a function of the sixteen Gains. The blue region of graphs indicates the Gains where the stimulus’ motion across the retina is in a direction that is directly opposite to eye motion. The green region indicates the Gains where the stimulus’ motion across the retina is in the same direction as eye motion. The top four rows show individual results from four subjects and the bottom row shows the averages across the five subjects, including 10003L whose data was shown in Fig. 2 C and D. Purple points indicate average diffusion constants for world motion, *D*_*WM*_, with SE of the mean bars. (**Left**) Yellow points indicate average diffusion constants for perceived motion, *D*_*PM*_, with SE of the mean bars. Black-hollow points indicate average *Jitter D*_*PM*_ with SE of the mean bars, given the standard deviation of error in Fig. A1.1. (**Right**) Orange points indicate average corrected diffusion constants for perceived motion, *cD*_*PM*_, with SE of the mean bars. *cD*_*PM*_ = *D*_*PM*_ − *Jitter D*_*PM*_.

**Figure A1.3:**
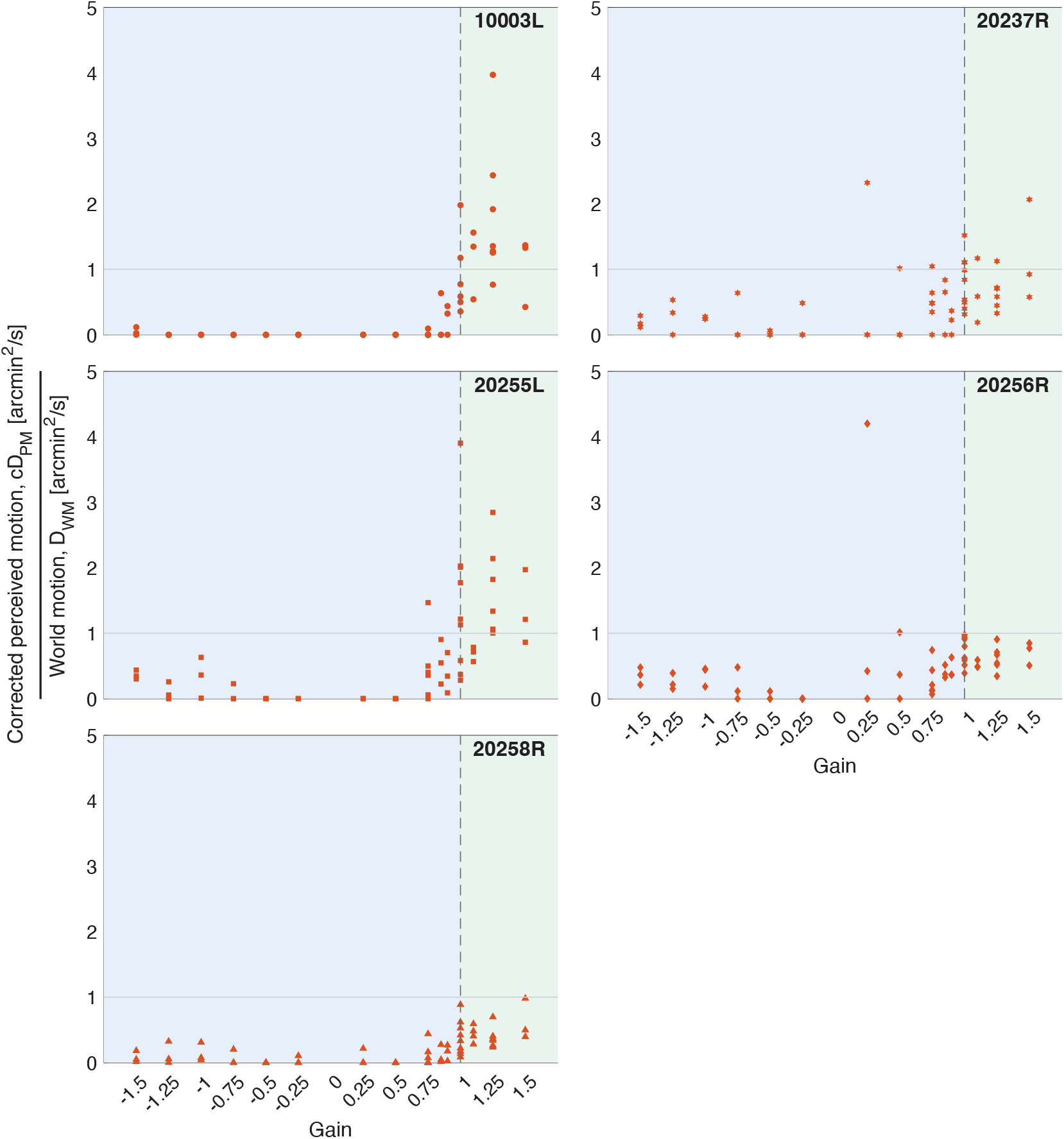
Experiment 1: Individual trial distribution of [average corrected diffusion constant for perceived motion]/[average diffusion constant for world motion], or *cD*_*PM*_ */D*_*WM*_, from five subjects as a function of the Gains. The blue regions indicate the Gains where the stimulus’ motion across the retina is in a direction that is directly opposite to eye motion. The green regions indicate the Gains where the stimulus’ motion across the retina is in the same direction as eye motion. Each point represents one trial. Note that ratios for Gain 0 were removed.

**Figure A1.4:**
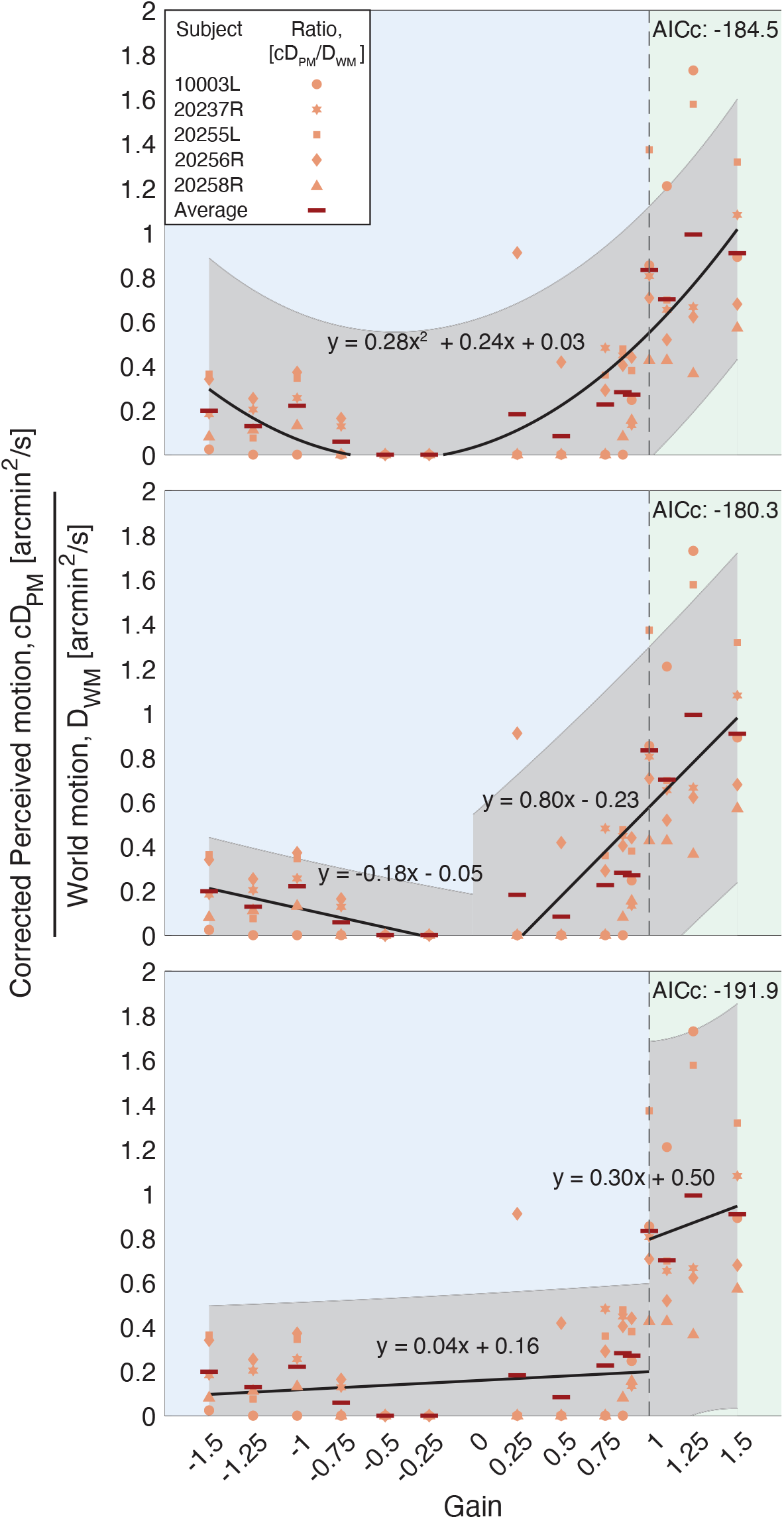
Experiment 1: Examples of linear and non-linear fits to the average ratios from five subjects. The blue regions indicate the Gains where the stimulus’ motion across the retina is in a direction that is directly opposite to eye motion. The green regions indicate the Gains where the stimulus’ motion across the retina is in the same direction as eye motion. Each point represents one subject and is the [average corrected diffusion constant for perceived motion]/[average diffusion constant for world motion], or *cD*_*PM*_ */D*_*WM*_. The dark red bars indicate the average ratio across the five subjects. The gray intervals indicate the 95% confidence bounds of the fits. The corrected Akaike information criterion, *AIC*_*c*_, values are labeled on the top right corner of each graph. **(Top Panel)** A single quadradic fit across all points. **(Middle Panel)** Two independent, two-parameter lines in slope-intercept form on either side of a breakpoint at Gain 0. **(Bottom Panel)** Two independent, two-parameter lines in slope-intercept form on either side of a breakpoint at Gain +1.

**Table A1.1:** Evaluation of different fits to the [average **corrected** diffusion constant for perceived motion]/[average diffusion constant for world motion], or **c***D*_*PM*_ */D*_*WM*_. Example fits are shown in Fig. 3B and Fig. A1.4 where *x* represents the Gain and *y*_*i*_ represents the *cD*_*PM*_ */D*_*WM*_. (A) We tested a quadradic, a cubic, and quartic function across all data points. We also tested multiple breakpoints ranging from Gain −1 to +1.1, and performed an independent linear fit to each side of the breakpoint. (B) We tested independent, single-parameter linear fits on either side of the break point. (C) We also tested independent, two-parameter linear fits in slope-intercept form on either side of the breakpoint.

| (a) One Continuous Fit |  |  |  |  |  |
| --- | --- | --- | --- | --- | --- |
| Fit | AICc | ΔAICc | Likelihood | w | evidenceRatio |
| Quadratic | -184.50 | 10.10 | 6.40E-03 | 4.97E-03 | 156.21 |
| Cubic | -183.68 | 10.92 | 4.24E-03 | 3.29E-03 | 235.64 |
| Quartic | -183.59 | 11.01 | 4.07E-03 | 3.16E-03 | 245.51 |

| (b) Two Horizontal Lines on Either Side of a Breakpoint |  |  |  |  |  |
| --- | --- | --- | --- | --- | --- |
| Breakpoint | AICc | ΔAICc | Likelihood | w | evidenceRatio |
| -1 | -128.99 | 65.61 | 5.66E-15 | 4.39E-15 | 1.77E+14 |
| -0.75 | -129.64 | 64.96 | 7.83E-15 | 6.08E-15 | 1.28E+14 |
| -0.5 | -132.85 | 61.75 | 3.91E-14 | 3.03E-14 | 2.56E+13 |
| -0.25 | -138.44 | 56.16 | 6.38E-13 | 4.95E-13 | 1.57E+12 |
| 0 | -146.28 | 48.32 | 3.22E-11 | 2.50E-11 | 3.10E+10 |
| 0.25 | -146.28 | 48.32 | 3.22E-11 | 2.50E-11 | 3.10E+10 |
| 0.5 | -150.58 | 44.02 | 2.76E-10 | 2.14E-10 | 3.62E+09 |
| 0.75 | -161.02 | 33.58 | 5.10E-08 | 3.96E-08 | 1.96E+07 |
| 0.85 | -168.69 | 25.91 | 2.36E-06 | 1.83E-06 | 4.24E+05 |
| 0.9 | -177.26 | 17.34 | 1.71E-04 | 1.33E-04 | 5.84E+03 |
| 1 | -194.60 | 0 | 1 | 0.78 | 1 |
| 1.1 | -168.01 | 26.59 | 1.68E-06 | 1.30E-06 | 5.96E+05 |

| (c) Two Lines in Slope-Intercept Form on Either Side of a Breakpoint |  |  |  |  |  |
| --- | --- | --- | --- | --- | --- |
| Breakpoint | AICc | ΔAICc | Likelihood | w | evidenceRatio |
| -1 | -158.53 | 36.07 | 1.47E-08 | 1.14E-08 | 6.81E+07 |
| -0.75 | -166.93 | 27.67 | 9.82E-07 | 7.62E-07 | 1.02E+06 |
| -0.5 | -171.49 | 23.11 | 9.59E-06 | 7.44E-06 | 1.04E+05 |
| -0.25 | -175.24 | 19.36 | 6.24E-05 | 4.84E-05 | 1.60E+04 |
| 0 | -180.32 | 14.28 | 7.92E-04 | 6.15E-04 | 1.26E+03 |
| 0.25 | -180.32 | 14.28 | 7.92E-04 | 6.15E-04 | 1.26E+03 |
| 0.5 | -183.62 | 10.98 | 4.12E-03 | 3.20E-03 | 242.61 |
| 0.75 | -183.81 | 10.79 | 4.54E-03 | 3.52E-03 | 220.30 |
| 0.85 | -182.82 | 11.78 | 2.77E-03 | 2.15E-03 | 361.50 |
| 0.9 | -183.29 | 11.31 | 3.50E-03 | 2.71E-03 | 286.10 |
| 1 | -191.89 | 2.72 | 0.26 | 0.20 | 3.89 |
| 1.1 | -171.02 | 23.58 | 7.59E-06 | 5.89E-06 | 1.32E+05 |

**Table A1.2:** Evaluation of different fits to the [average diffusion constant for perceived motion]/[average diffusion constant for world motion], or *D*_*PM*_ */D*_*WM*_. (Note that the *D*_*PM*_ are the original, uncorrected values.) (A) We tested a quadradic, a cubic, and quartic function across all data points. We also tested multiple breakpoints ranging from Gain −1 to +1.1, and performed an independent linear fit to each side of the breakpoint. (B) We tested independent, single-parameter linear fits on either side of the break point. (C) We also tested independent, two-parameter linear fits in slope-intercept form on either side of the breakpoint.

| (a) One Continuous Fit |  |  |  |  |  |
| --- | --- | --- | --- | --- | --- |
| Fit | AICc | ΔAICc | Likelihood | w | evidenceRatio |
| Quadratic | -153.90 | 6.22 | 0.044 | 0.026 | 22.47 |
| Cubic | -152.47 | 7.65 | 0.022 | 0.013 | 45.87 |
| Quartic | -150.22 | 9.90 | 7.08E-03 | 4.19E-03 | 141.18 |

| (b) Two Horizontal Lines on Either Side of a Breakpoint |  |  |  |  |  |
| --- | --- | --- | --- | --- | --- |
| Breakpoint | AICc | ΔAICc | Likelihood | w | evidenceRatio |
| -1 | -128.54 | 31.59 | 1.38E-07 | 8.19E-08 | 7.22E+06 |
| -0.75 | -129.51 | 30.61 | 2.26E-07 | 1.33E-07 | 4.43E+06 |
| -0.5 | -133.26 | 26.86 | 1.47E-06 | 8.70E-07 | 6.80E+05 |
| -0.25 | -140.43 | 19.69 | 5.30E-05 | 3.10E-05 | 1.89E+04 |
| 0 | -140.85 | 19.27 | 6.54E-05 | 3.87E-05 | 1.53E+04 |
| 0.25 | -140.85 | 19.27 | 6.54E-05 | 3.87E-05 | 1.53E+04 |
| 0.5 | -137.78 | 22.34 | 1.41E-05 | 8.32E-06 | 7.11E+04 |
| 0.75 | -140.64 | 19.48 | 5.89E-05 | 3.48E-05 | 1.70E+04 |
| 0.85 | -145.83 | 14.29 | 7.88E-04 | 4.66E-04 | 1.27E+03 |
| 0.9 | -150.39 | 9.73 | 7.70E-03 | 4.55E-03 | 129.91 |
| 1 | -160.12 | 0 | 1 | 0.59 | 1 |
| 1.1 | -147.84 | 12.29 | 2.15E-03 | 1.27E-03 | 465.72 |

| (c) Two Lines in Slope-Intercept Form on Either Side of a Breakpoint |  |  |  |  |  |
| --- | --- | --- | --- | --- | --- |
| Breakpoint | AICc | ΔAICc | Likelihood | w | evidenceRatio |
| -1 | -144.61 | 15.51 | 4.28E-04 | 2.53E-04 | 2.34E+03 |
| -0.75 | -146.58 | 13.54 | 1.15E-03 | 6.77E-04 | 873.07 |
| -0.5 | -147.01 | 13.11 | 1.42E-03 | 8.42E-04 | 702.31 |
| -0.25 | -146.84 | 13.28 | 1.30E-03 | 7.71E-04 | 766.88 |
| 0 | -147.77 | 12.36 | 2.07E-03 | 1.23E-03 | 482.13 |
| 0.25 | -147.77 | 12.36 | 2.07E-03 | 1.23E-03 | 482.13 |
| 0.5 | -152.65 | 7.47 | 0.024 | 0.014 | 41.88 |
| 0.75 | -153.34 | 6.78 | 0.034 | 0.020 | 29.65 |
| 0.85 | -153.02 | 7.10 | 0.029 | 0.017 | 34.82 |
| 0.9 | -153.71 | 6.41 | 0.040 | 0.024 | 24.70 |
| 1 | -158.57 | 1.55 | 0.46 | 0.27 | 2.17 |
| 1.1 | -151.02 | 9.10 | 0.011 | 6.24E-03 | 94.83 |

**Figure A1.5:**
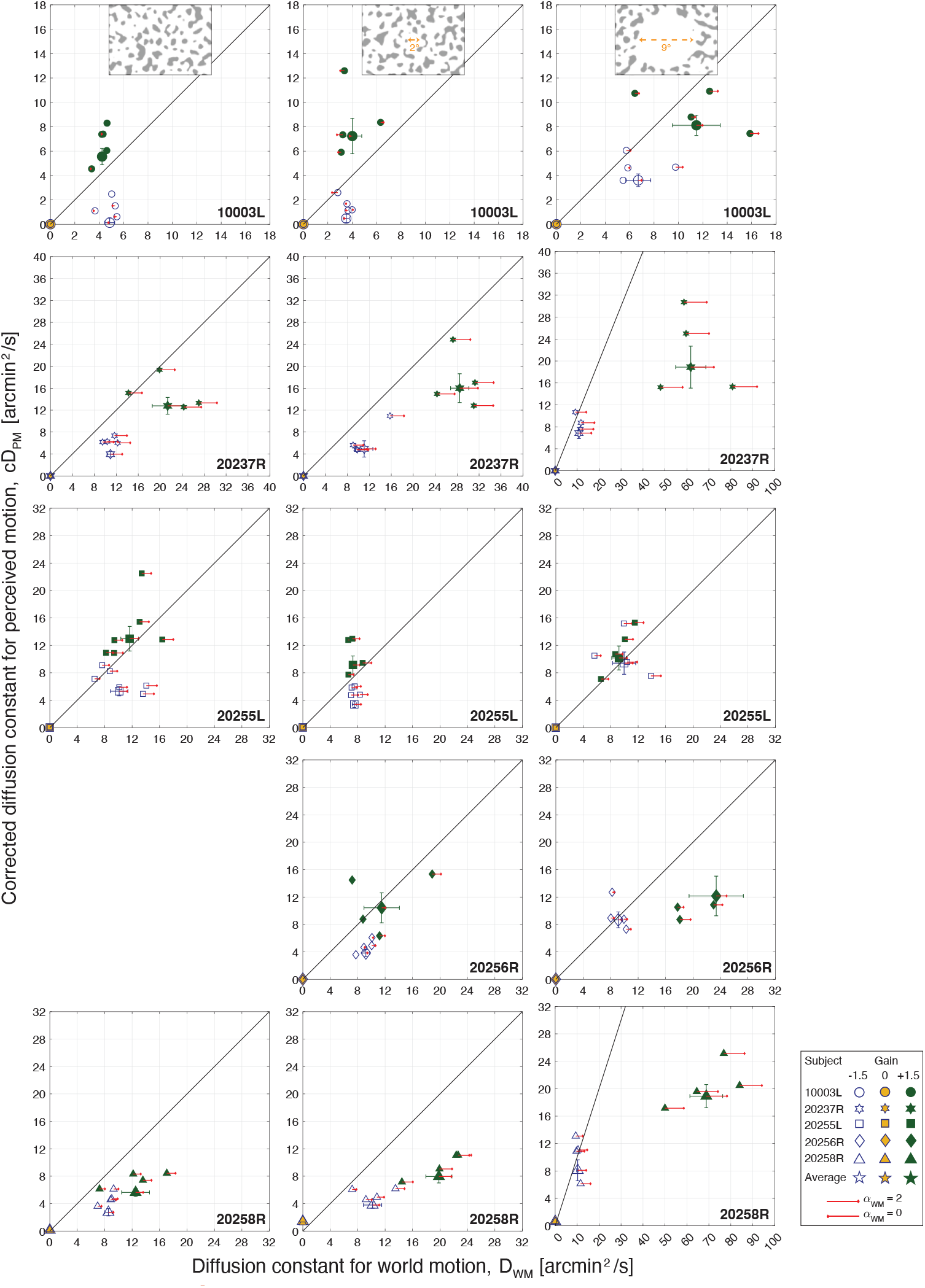
Experiment 2: Corrected diffusion constants for perceived motion (*cD*_*PM*_) versus diffusion constants for world motion (*D*_*WM*_) from five subjects. Experiments were tested under three background conditions: The patterns extended over the full 17° field of view (**Left**), or surrounded a 2°-diameter white circular opening (**Middle**), or surrounded a 9°-diameter white circular opening (**Right**). Background conditions are indicated by labels on the top row. Note that subject 20256R was not tested under the no-white-circle condition. Each small symbol represents one trial: for Gain −1.5 stimuli (blue open symbols), Gain 0 stimuli (yellow filled symbols), and Gain +1.5 stimuli (green filled symbols). The large stars are the averages with SE of the mean bars. The y-axes are different to account for individual differences in eye motion and consequent retina-contingent stimulus’ world motion (*D*_*WM*_). The red arrows show the extent to which the eye motion, and consequent retina contingent stimulus’ world motion (*α*_*WM*_), deviated from Brownian. Arrows pointing Right indicate persistence (*α*_*WM*_ *>* 1), arrows pointing Left indicate antipersistence (*α*_*WM*_ *<* 1), and no arrow means that the motion was Brownian (*α*_*WM*_ = 1 +*/*− 0.02). Longer arrows correspond to higher deviations from Brownian motion. The arrow length in the legend indicates pure persistence (*α*_*WM*_ = 2, straight line trajectory at constant velocity) if pointing Right or pure antipersistence (*α*_*WM*_ = 0, oscillatory motion) if pointing Left.

**Figure A1.6:**
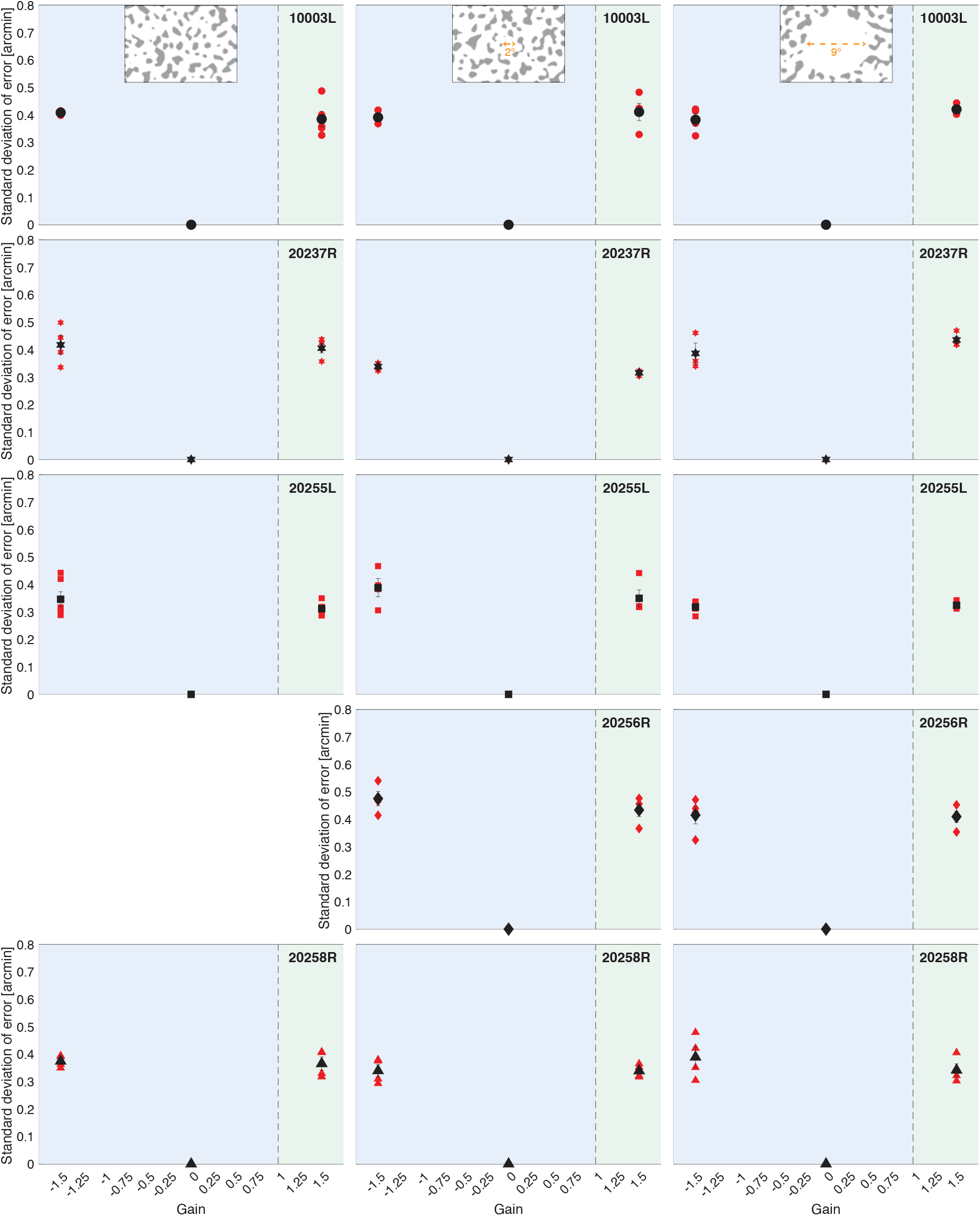
Experiment 2: The magnitude of stimulus delivery error for each Gain from five subjects. The blue regions indicate the Gains where the stimulus’ motion across the retina is in a direction that is directly opposite to eye motion. The green region indicates the Gains where the stimulus’ motion across the retina is in the same direction as eye motion. Experiments were tested under three background conditions: The patterns extended over the full 17° field of view (**Left**), or surrounded a 2°-diameter white circular opening (**Middle**), or surrounded a 9°-diameter white circular opening (**Right**). Background conditions are indicated by labels on the top row. For each trial, we computed the the distances in x and y between the ideal retina-contingent stimulus motion trace and the measured retina-contingent stimulus motion trace. The red points indicate the standard deviation of error from one trial – this is the standard deviation across the distances in x and y from all frames from all videos within a single trial. The black points indicate the average standard deviation of error across all trials, with SE of the mean bars.

**Figure A1.7:**
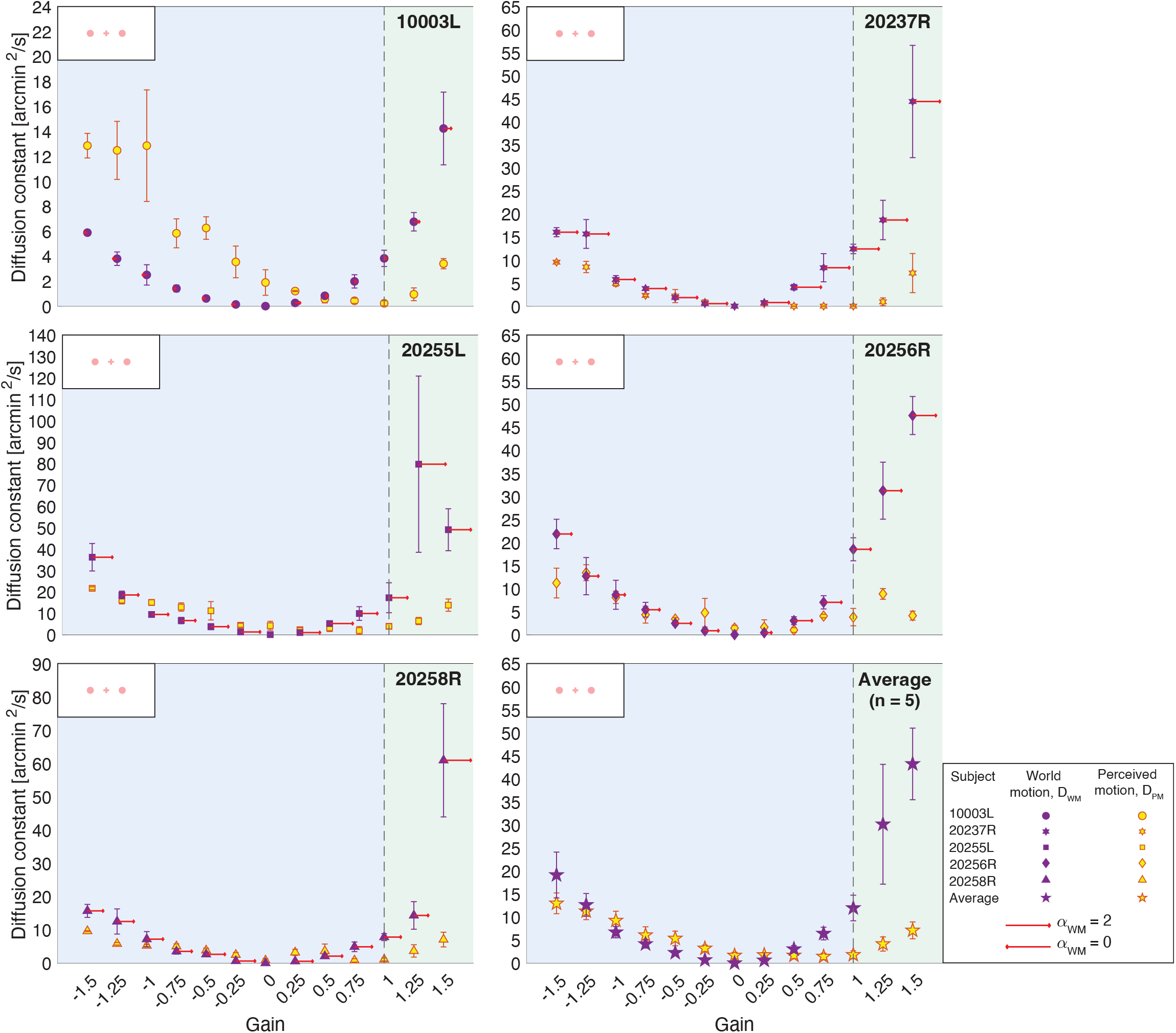
Control Experiment: Results from experiments tested under background-absent “Ganzfeld” conditions, indicated by labels on the top left corner of each graph. The blue regions indicate the Gains where the stimulus’ motion across the retina is in a direction that is directly opposite to eye motion. The green regions indicate the Gains where the stimulus’ motion across the retina is in the same direction as eye motion. Note that the stimuli were presented for 750-ms for this condition. Average diffusion constants are plotted as a function of the Gains from five subjects. The orange symbols represent diffusion constants for perceived motion (*D*_*PM*_) and the purple symbols represent diffusion constants for world motion (*D*_*WM*_). (Note that the *D*_*PM*_ are the original, uncorrected values.) The y-axes are different to account for individual differences in eye motion and consequent retina-contingent stimulus’ world motion (*D*_*WM*_). Each symbol is the respective average across all trials, with SE of the mean bars. The bottom right graph shows the averages across the five subjects, with SE of the mean bars. The red arrows show the extent to which the eye motion, and consequent retina-contingent stimulus’ world motion (*α*_*WM*_), deviated from Brownian. Arrows pointing right indicate persistence (*α*_*WM*_ *>* 1), arrows pointing left indicate antipersistence (*α*_*WM*_ *<* 1), and no arrow means that the motion was Brownian (*α*_*WM*_ = 1 +/− 0.02). Longer arrows correspond to higher deviations from Brownian motion. The arrow length in the legend indicates pure persistence (*α*_*WM*_ = 2, straight line trajectory at constant velocity) if pointing right or pure antipersistence (*α*_*WM*_ = 0, oscillatory motion) if pointing left.

## Appendix 2: Comparison with Previous Work

In Arathorn et al. (2013) and D’Angelo et al. (2024), under background-present conditions, stimuli moving in the same direction as eye motion (Gains ≥ +1) appeared readily in motion. For Gains ≥ +1, we would expect that the motion the subjects perceived would be equal to the motion of the stimuli in the world. Arathorn et al. (2013) did not quantify the magnitude of eye motion or the retina contingent stimulus’ world motion so we cannot make any claims on whether the subjects veridically perceived the world motion of the retina-contingent stimuli. But D’Angelo et al. (2024) computed the diffusion constant and *α* of the eye motion (*D*_*EM*_ and *α*_*EM*_) and retina-contingent stimulus’ world motion (*D*_*WM*_ and *α*_*WM*_); the same method is used in this study, so we can compare the two studies, shown in Fig. A2.1A.

Fig. A2.1A plots the average diffusion constants for perceived motion (*D*_*PM*_) versus diffusion constants for world motion (*D*_*WM*_). (Note that the *D*_*PM*_ are the original, uncorrected values.) Results were collected from five subjects for this current paper and six subjects for D’Angelo et al. (2024), indicated by light and dark colors, respectively. For both studies, experiments were tested under background-present conditions: the background was filled with a blurred and binarized 1/f noise pattern with a 2°-diameter central white circle with a surrounding black ring that overlayed the 680-nm stimuli, shown in Fig. 1A. The small symbols represent each subject’s average perceptual match for Gain −1.5 stimuli (light- and dark-blue open symbols), Gain 0 stimuli (light- and dark-yellow filled symbols), and Gain +1.5 stimuli (light- and dark-green filled symbols). The 1:1 indicates veridical perception of motion.

In this study, for Gains +1.5 (light green data points in Fig. A2.1A), three subjects overestimated the motion (10003L, 20237R, and 20255L) and two subjects underestimated the motion (20256R and 20258R). As a result, the average perceived motion across subjects (large light-green star in Fig. A2.1A) was approximately equal to the world motion of the stimuli. In D’Angelo et al. (2024), for Gains +1.5 (dark green data points in Fig. A2.1A), five subjects underestimated the motion and only one subject (10003L) reported motion that was similar to the world motion. As a result, the average perceived motion across subjects (large dark-green star in Fig. A2.1A) was below the world motion of the stimuli.

We will discuss three factors that could contribute to this:

### 1. Inter- and intra-subject variability

Participants were different between studies. We may have by random chance recruited subjects who were naturally biased to underestimate motion in D’Angelo et al. (2024) (inter-subject variability). It is important to mention that in each study, no specific criteria were given to the subjects to make the match, so the subjects who were recruited for both studies may have changed criteria between the studies (intra-subject variability). As shown in Fig. A2.1A, for Gain +1.5 stimuli, 10003L (circle) performed similarly between both studies, but 20237R (hexagram) and 20256R (diamond) perceived more motion in this current paper compared to D’Angelo et al. (2024).

### 2. Eccentricity

We presented stimuli 0.43° away from the fovea in this study; however, stimuli were delivered 2° away from the fovea in D’Angelo et al. (2024). Motion perception thresholds have been shown to increase with eccentricity (Bower, Bian, & Andersen, 2012; McKee & Nakayama, 1984). Interestingly, the perceived motion of Gain −1.5 stimuli are also higher in this current paper (light- vs dark-blue in Fig. A2.1A).

### 3. α for world motion, α_WM_

We discussed in D’Angelo et al. (2024) that a higher *α* leads to a higher diffusion constant (D), given the same average step length. (“Step length” refers to the standard deviation of the randomized step lengths within a trajectory). In other words, for a subject with persistent eye motion, the subsequent retina-contingent stimulus has a high *α*_*WM*_ and a higher diffusion constant compared to a trajectory with the same average step length but with an *α* equal to 1. Since the subject always compares a Brownian random walk stimulus to their potentially highly persistent retina-contingent stimulus motions, this could lead to high diffusion constants for world motion (*D*_*WM*_), leading to smaller *D*_*PM*_ */D*_*WM*_ that convey underestimated values of motion perceptions. This was especially relevant in the Ganzfeld condition, where the *α*_*WM*_ ‘s were as high as 1.7. (Note that an *α* of 2 is maximum persistence, a straight line trajectory at constant velocity.) Subjects expressed no frustration when asked to match a persistent trajectory to a Brownian trajectory. Reported in D’Angelo et al. (2024), subjects perform poorly at discriminating a highly persistent motion (e.g. 1.3 *< α <* 1.7) with a high diffusion constant from a less persistent motion (e.g. 1 *< α <* 1.3) with a low diffusion constant.

But, under background-present conditions, the *α*_*WM*_ ‘s are similar between D’Angelo et al. (2024) and this current paper (indicated by red arrows in Figure A2.1A). *α*_*WM*_ cannot fully explain the motion perception differences between the two studies. We plotted the [average diffusion constant for perceived motion]/[average diffusion constant for world motion], or *D*_*PM*_ */D*_*WM*_, from Fig. 3B as a function of each subject’s *α*_*WM*_, for four Gains: +1, +1.1, +1.25, and +1.5. The respective colors go from light to dark green, shown in Fig. A2.1B. Using the same procedure from D’Angelo et al. (2024), we modeled how the ratios would vary with *α*_*WM*_ under a specific condition where the subject equated the mean square displacement (MSD) between the motion of the random walk stimulus over a specific time interval. In the model, the random walk stimulus’ *α*, which represents the *α*_*PM*_, is equal to one (Brownian motion), while the retina-contingent stimulus’ *α*_*WM*_ ranges from antipersistent (0.9 ≤ *α*_*WM*_ *<* 1), to Brownian (*α*_*WM*_ = 1), to persistent (1 *< α*_*WM*_ ≤ 1.8). By a least-squares procedure, we determined that a time interval of 28 frames best predicted the measured data. The model is represented by a solid curve in Fig. A2.1B.

In D’Angelo et al. (2024), computing the MSD over a time interval of 2 frames best predicted the measured data. This analysis suggests that subjects make motion judgments for perifoveal stimuli over longer time intervals (14x longer) in this study, compared to stimuli presented at greater eccentricities. This could also be due to the stimuli being presented side-by-side instead of over intervals of time. Further work could evaluate each subjects’ specific time interval over which they make motion judgments.

**Figure A2.1:**
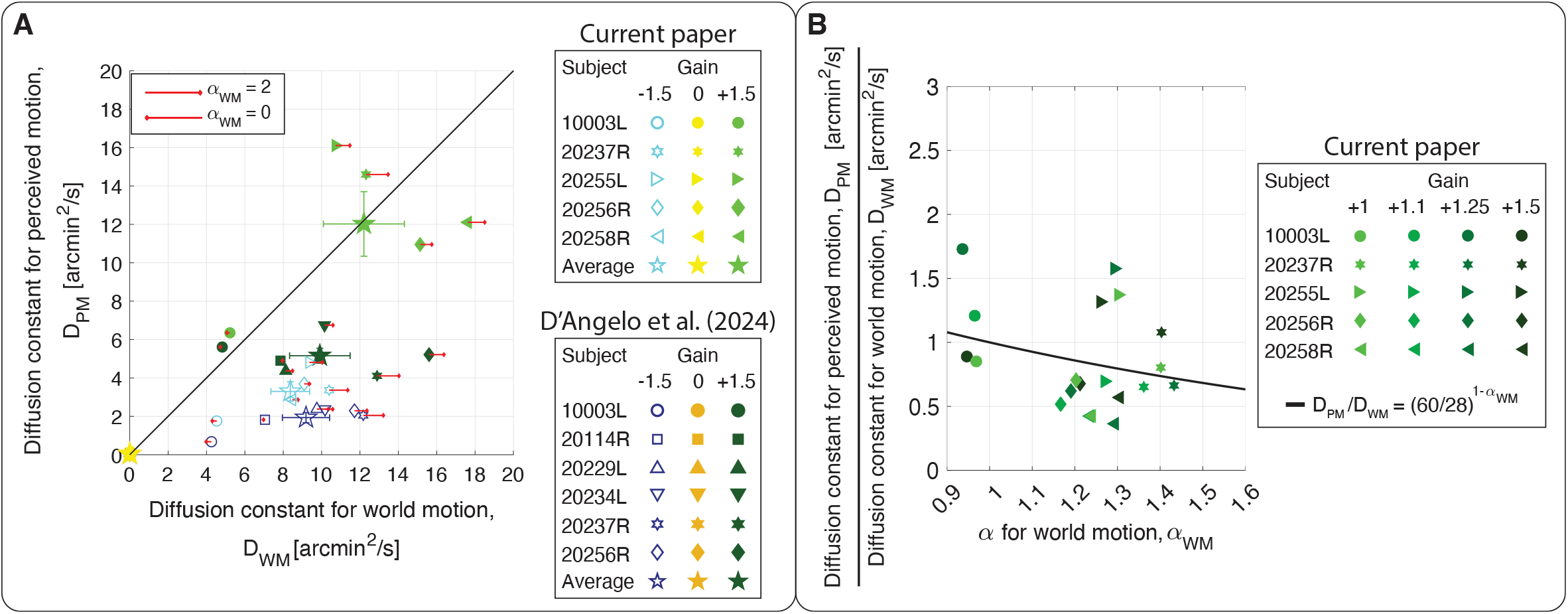
Comparing results from this paper to D’Angelo et al. (2024). Average diffusion constants for perceived motion (*D*_*PM*_) versus diffusion constants for world motion (*D*_*WM*_). (Note that the *D*_*PM*_ are the original, uncorrected values.) Results were collected from five subjects for this current paper and six subjects for D’Angelo et al. (2024), indicated by light and dark colors, respectively. All experiments were tested under background-present conditions. The small symbols represent each subject’s average perceptual match for Gain −1.5 stimuli (light- and dark-blue open symbols), Gain 0 stimuli (light- and dark-yellow filled symbols), and Gain +1.5 stimuli (light- and dark-green filled symbols). The large stars are the group averages with SE of the mean bars. The red arrows show the extent to which the eye motion, and consequent retina-contingent stimulus’ world motion (*α*_*WM*_), deviated from Brownian. Arrows pointing right indicate persistence (*α*_*WM*_ *>* 1), arrows pointing left indicate antipersistence (*α*_*WM*_ *<* 1), and no arrow means that the motion was Brownian (*α*_*WM*_ = 1 +/− 0.02). Longer arrows correspond to higher deviations from Brownian motion. The arrow length in the legend indicates pure persistence (*α*_*WM*_ = 2, straight line trajectory at constant velocity) if pointing right or pure antipersistence (*α*_*WM*_ = 0, oscillatory motion) if pointing left. (B) Computed ratios from five subjects plotted as a function of each subject’s *α* for world motion, *α*_*WM*_. The symbols represent each subject’s [average diffusion constant for perceived motion]/[average diffusion constant for world motion], or *D*_*PM*_ */D*_*WM*_. (Note that the *D*_*PM*_ are the original, uncorrected values.) Data are from four Gains: +1, +1.1, +1.25, and +1.5, and the respective colors go from light to dark green. The black curve predicts the *D*_*PM*_ */D*_*WM*_ versus *α*_*WM*_, considering that the subject matched the mean squared displacements of the random walk stimulus with the retina-contingent stimulus over a time interval of 28 frames.

